# Dual cholinergic and serotonergic excitatory pathways mediate oxygen sensing in the zebrafish gill

**DOI:** 10.1101/2025.05.15.654070

**Authors:** Maddison Reed, Jan A. Mennigen, Michael G. Jonz

## Abstract

The evolution of oxygen sensing included a transition from a diffuse distribution of respiratory chemoreceptors in the gills of water-breathing vertebrates to chemoreceptor clusters confined to the pulmonary epithelium and carotid body in air-breathers. Since the excitatory neurotransmitters mediating oxygen sensing in anamniotes have never been confirmed, the origins of oxygen sensing in vertebrates have remained controversial. In gills isolated from Tg(*elavl3*:GCaMP6s) zebrafish expressing a genetically-encoded reporter of intracellular Ca^2+^ concentration ([Ca^2+^]_i_), we demonstrate that acetylcholine (ACh) and nicotine induced a dose-dependent increase in [Ca^2+^]_i_ in postsynaptic sensory neurons innervating oxygen-chemoreceptive neuroepithelial cells (NECs). Hypoxic stimulation of NECs evoked a similar rise in neuronal [Ca^2+^]_i_ that was abolished by nicotinic antagonist, hexamethonium. Using immunohistochemistry and RT-qPCR, we identified a novel population of ACh-containing NECs associated with sensory neurons expressing the α2 subunit of nicotinic ACh receptors. *In vivo* whole-larva Ca^2+^ imaging showed that cholinergic and hypoxic activation of the gills generated Ca^2+^ activity in neurons of vagal sensory ganglia with time-dependent characteristics of neurotransmission towards the hindbrain. We identified a second source of hypoxic activity in vagal sensory ganglia operating exclusively through 5-HT_3_ receptors and dependent upon vesicular monoamine transport (VMAT2) in the gill. We traced expression of 5-HT_3_ receptors to nerve terminals surrounding a separate population of serotonergic VMAT2-positive NECs. Our investigation reveals independent cholinergic and serotonergic autonomic pathways of oxygen sensing in zebrafish and provides the first physiological evidence that gill chemoreceptors may be homologues of both pulmonary and carotid body chemoreceptors in mammals.

**Significance statement:** The excitatory neurotransmitters mediating oxygen sensing in anamniotes have never been confirmed. Thus, the origins of oxygen sensing in vertebrates have remained controversial. In transgenic zebrafish expressing a genetically-encoded reporter of intracellular Ca^2+^ concentration, we identified two independent pathways of oxygen sensing in the gill: one involving acetylcholine and interneurons intrinsic to the gill, and the other via serotonin acting directly upon ganglionic neurons. Both pathways resulted in excitation of vagal sensory ganglia that receive hypoxic inputs from the gills and innervate the hindbrain. We argue that gill chemoreceptors are homologues of both pulmonary and carotid body chemoreceptors in mammals.

## Introduction

Oxygen sensing and the physiological responses to hypoxia are initiated by specialized respiratory chemoreceptors. The organization of chemoreceptors controlling respiratory drive underwent an evolutionary transition from a diffuse distribution in the gills of water-breathing anamniotes to a localized arrangement of chemoreceptors in air-breathing vertebrates where, in mammals, chemoreceptors are found as neuroepithelial bodies (NEBs) of the pulmonary epithelium and clusters of type I cells in the carotid body (Milsom and Burleson, 2007; Kumar and Prabhakar, 2012; Cutz et al., 2013; Hockman et al., 2017). The gills of aquatic vertebrates, such as fish, therefore serve as important models for understanding how oxygen sensing evolved in vertebrates.

The gill is a primary site of gas exchange and detection of hypoxia. Gill neuroepithelial cells (NECs) are the major oxygen-sensing chemoreceptors in aquatic vertebrates and share morphological and physiological features with those of mammals (Dunel-Erb et al., 1982; Porteus et al., 2012; Reed and Jonz, 2022; Leonard et al., 2024). In zebrafish (*Danio rerio*), NECs respond to hypoxia through inhibition of background K^+^ channels, membrane depolarization, elevation of intracellular Ca^2+^ concentration ([Ca^2+^]_i_) and subsequent neurotransmitter release (Jonz et al., 2004; Zachar et al., 2017a; Reed and Jonz, 2025). This process activates sensory pathways in the gill, ultimately resulting in autonomic reflexes, such as hyperventilation, that improve oxygen uptake (Perry et al., 2009). Gill NECs in zebrafish are innervated and become functional during the larval period, when they begin to display adult-like responses to hypoxia (Jonz and Nurse, 2005; Pan et al., 2019). Multiple populations of NECs have been identified in zebrafish, all of which express the synaptic vesicle protein-2 (SV2) and may contain serotonin (5-hydroxytryptamine, 5-HT), the vesicular acetylcholine transporter (VAChT) or other neurotransmitters (Jonz and Nurse, 2003; Zachar et al., 2017b). Dopamine was recently identified as a presynaptic modulator of the NEC response to hypoxia (Reed and Jonz, 2025), but since the excitatory neurotransmitters responsible for hypoxic signaling in the gill have not yet been identified, the homology between gill and mammalian chemoreceptors has remained controversial.

Fish gills are innervated by branches of the glossopharyngeal (IX) and vagus (X) nerves and NECs receive innervation from extrinsic neurons originating in the hindbrain and vagal sensory ganglia, and from neurons intrinsic to the gills (Donald, 1987; Bailly et al., 1992; Sundin and Nilsson, 2002; Jonz and Nurse, 2003). One source of intrinsic innervation arises from a series of sensory neurons resembling a "chain" (ChNs; Jonz and Nurse, 2003), whose cell bodies are distributed along the gill filament artery and send projections to innervate NECs (Jonz and Nurse, 2003; Reed et al., 2024). Recent studies using transgenic Tg(*elavl3*:GCaMP6s) zebrafish, which express a genetically-encoded Ca^2+^ reporter under the control of a pan-neuronal promoter, have provided insights into the functional aspects of the NEC-ChN synapse (Reed and Jonz, 2025). In these animals, both NECs and postsynaptic ChNs display a hypoxia-induced increase in [Ca^2+^]_i_, but the ChN response is dependent upon functional synaptic contact with oxygen-sensitive NECs.

Serotonin, the most abundant neurotransmitter in gill NECs, and acetylcholine (ACh), have both been suggested as excitatory neurotransmitters (Jonz and Nurse, 2003; Porteus et al., 2012; Zachar et al., 2017b). Furthermore, single-cell RNA-sequencing in zebrafish has provided deeper insights and revealed that NECs and gill neurons express genes associated with chemosensory transduction of hypoxic signals, such as genes encoding 5-HT and nicotinic Ach receptors (nAChRs), many of which are shared with mammalian chemoreceptors (Pan et al., 2022).

The goal of the present study was to uncover the excitatory pathways in the gill involved in hypoxia signaling. Using a combination of intracellular Ca^2+^ imaging, immunohistochemistry and gene expression analysis, we provide evidence for ACh as a key excitatory neurotransmitter acting through nAChRs at the NEC-ChN synapse. We further identify an additional serotonergic pathway for hypoxia signaling acting through gill 5-HT_3_ receptors that operates independently of ChNs. Stimulation of both serotonergic and cholinergic receptor pathways results in activation of the vagal sensory ganglia involved in control of ventilation.

## Methods

### Animal ethics statement

All wild-type and transgenic zebrafish were bred and maintained it the Laboratory for the Physiology and Genetics of Aquatic Organisms, University of Ottawa. Zebrafish were kept at 28°C on a 14:10-h light:dark cycle (Westerfield, 2007). Embryos and larvae were reared on a diet of rotifers and Gemma 75 (Skretting Canada, St. Andrews, NB). Juveniles were fed Artemia and Gemma 150-300 (Skretting) and adults were fed Adult Zebrafish Diet (Ziegler Feeds, Gardners, PA, USA). Adult zebrafish were euthanized by concussion and terminated by decapitation. Larval zebrafish were euthanized by immersion in an ice bath for 20 min. All procedures for animal use and euthanasia were carried out in accordance with institutional guidelines according to protocol BL-3666, and guidelines provided by the Canadian Council on Animal Care.

### Relative [Ca^2+^]_i_ measurements

#### Ex vivo gill recording

In this study we used transgenic zebrafish Tg(*elavl3*:GCaMP6s) expressing the genetically encoded Ca^2+^ indicator, GCaMP6s, under the pan-neuronal promotor, *elavl3* (Dunn et al., 2016; Reed and Jonz, 2025). GCaMP6s contains the green fluorescent protein (GFP) as part of its structure. Whole gill baskets were removed and separated into individual gill arches and immersed in extracellular solution containing (mM): 120 NaCl, 5 KCl, 2.5 CaCl_2_, 2 MgCl_2_, 10 HEPES, 10 glucose at pH 7.8. Isolated intact gill arches from Tg(*elavl3*:GCaMP6s) zebrafish were secured in a Petri dish using a metal tissue anchor (cat. no. 640251, Warner Instruments, Holliston, MA, USA) and continuously superfused with extracellular solution at pH 7.8. GFP-fluorescing cells were observed using a 40× water-immersion objective (Nikon, Tokyo, Japan). GCaMP-containing ChNs were identified by their distinct morphology and presence of GFP, as previously described (Reed and Jonz, 2025). Using a Lambda DG-5 wavelength changer (Sutter Instruments, Novato, CA, USA), the preparation was exposed to 490 nm excitation light for 600 ms at a sampling frequency of 1 s^-1^. Images were captured with a CCD camera (QImaging, Surrey, BC, Canada) and fluorescence intensity was recorded with NIS Elements software (Nikon). For each recording, the baseline (control) fluorescence was calculated as the average fluorescence intensity of a cell for the first 30 s in normoxia. All fluorescence values were divided by the baseline to evaluate changes in fluorescence intensity (F/F_0_) over time throughout a single recording.

#### In vivo recording from vagal sensory ganglia

Larval nerve recordings were carried out with whole, anesthetized animals from the Tg(*elavl3*:GCaMP6s) line aged 14-21 days post-fertilization. Animals were anesthetized with 0.01 mg/ml tricaine and held in place under a metal tissue anchor. Nodose and epibranchial ganglia (which together comprise the vagal sensory ganglia) were identified by position relative to the eye, otolith, gills and hindbrain. Recording regions were set to encompass the entire visible cluster of neurons associated with these ganglia (see Fig. 7A-C). For simultaneous recordings of epibranchial and nodose ganglia, sampling frequency was increased to 4 s^-1^ to more accurately observe the delay (latency) in activation time between these structures. Latency was defined as the difference between nodose activation time and epibranchial activation time (Time_nodose_ – Time_epibranchial_). Onset of the [Ca^2+^]_i_ response was characterized as the time where relative fluorescence first doubled (F/F_0_=2). Larval recording experiments were otherwise performed as described for whole-gill recording, including solutions, equipment setup and calculation of fluorescence intensity. All drugs were introduced into the recording chamber by perfusion in extracellular solution.

### Solutions and drug treatments

High K^+^ extracellular solution was prepared with 90 mM NaCl, 35 mM KCl, 2.5 mM CaCl_2_, 2 mM MgCl_2_, 10 mM HEPES, 10 mM glucose. Zero Ca^2+^ extracellular solution was prepared with 120 mM NaCl, 5 mM KCl, 4.5 mM MgCl_2_, 10 mM HEPES, 10 mM glucose, and 1 mM EGTA. pH of all solutions was kept at 7.8. 100% N_2_ was bubbled through an air stone into solution reservoirs to create hypoxic solutions with a PO_2_ of approximately 25 mmHg. All control solutions were bubbled with compressed air for the same duration of time.

Drugs were introduced into the recording chamber in extracellular solution. Acetylcholine (cat. no. A6625, Sigma-Aldrich, Oakville, ON, Canada) and nicotine (cat. no. N5260, Sigma-Aldrich) were used to activate nAChRs, and hexamethonium (cat. no. H2138, Sigma-Aldrich) was used as a nAChR blocker. To target 5-HT receptors, 5-HT (cat. no. H9523, Sigma-Aldrich), the 5-HT_3_ receptor agonist, 1-phenylbiguanide (cat. no. 164216, Sigma-Aldrich), and the 5-HT_3_ receptor antagonist, MDL 72222 (cat. no. 0640, Tocris, Toronto, ON, Canada), were tested. To deplete endogenous 5-HT from gill NECs, larvae were incubated in the VMAT2 blocker, tetrabenazine (cat. no. T2592, Sigma-Aldrich), for 20 min prior to hypoxia exposure. All drugs were first dissolved in dimethyl sulfoxide (DMSO) to produce a final DMSO concentration of <0.1%. At this concentration, DMSO had no effect on Ca^2+^ baseline and did not produce changes in fluorescence intensity (relative [Ca^2+^]_i_) in the isolated gill or the whole animal preparations.

### Immunohistochemistry

Techniques for tissue extraction and immunolabeling were carried out as previously described (Jonz and Nurse, 2003). Whole gill baskets were removed and immersed in phosphate buffered solution (PBS) containing (mM): NaCl 137, Na_2_HPO_4_ 15.2, KCl 2.7, and KH_2_PO_4_ 1.5 at pH 7.8. Gill baskets were fixed by immersion in 4% paraformaldehyde in PBS overnight at 4°C. Tissues were removed and rinsed in PBS three times at 3 min before permeabilization for 24 h at 4°C. Permeabilizing solution (PBS-TX) contained 0.5-2% Triton X-100 in PBS (pH 7.8). After 3 rinses in PBS, gill baskets were then separated into individual arches. Gill arches were incubated in primary antibodies for 24 h at 4°C, rinsed with PBS three times at 3 min, and immersed in secondary antibodies for 1 h at room temperature in darkness.

Acetylcholine-positive cells were identified using polyclonal anti-ACh raised in rabbit (RRID: AB_958833; cat. no. NB100-64656; Bio-Techne Canada, Toronto, ON). Pre-adsorption of ACh (cat. no. A6625, Sigma-Aldrich) with the anti-ACh antibodies blocked all immunolabeling in gill tissue. The nAChR α2 subunit was identified using monoclonal anti-CHRNA2 raised in mouse (RRID: AB_2787307; cat. no. MA5-31683, Invitrogen). Both antibodies were used at 1:100 and visualized with goat anti-rabbit secondary antibodies conjugated with fluorescein isothiocyanate (FITC, 1:50, cat. no. 111-095-003, Cedarlane, Burlington, ON, Canada). To control for the specificity of α2 antibodies in our preparation we used the specific control peptide corresponding to the immunogen sequence (DLEQMERTVDLKD, Alomone, BLP-NC002). Pre-adsorption of the control peptide with the α2 antibodies blocked all immunolabeling in gill tissue.

All NECs were identified using monoclonal SV2 raised in mouse (RRID: AB_2315387; Developmental Studies Hybridoma Bank, University of Iowa, IA, USA) at 1:100 and visualized by goat anti-mouse secondary antibodies conjugated with Alexa 594 at 1:100 (cat. no. A11005, Invitrogen, Burlington, ON, Canada). Monoclonal zn-12 raised in mouse (RRID: AB_531908; Developmental Studies Hybridoma Bank) was used to visualize gill neurons. zn-12 targets membrane fractions from adult zebrafish CNS and recognizes an HNK-1-like epitope (manufacturer specifications). zn-12 was used at 1:100 and targeted by goat anti-mouse secondary antibodies conjugated with Alexa 594 at 1:100 (cat. no. A11005; Invitrogen). Labeling of SV2 and zn-12 in the zebrafish gill has been previously characterized (Jonz and Nurse, 2003).

Serotonin-positive NECs were identified using the transgenic ET(*vmat2:GFP*) zebrafish obtained from the Becker Laboratory at the University of Edinburgh, as previously described (Pan et al., 2021). VMAT2 is an integral membrane protein that mediates storage of monoamines, such as 5-HT, into synaptic vesicles. ET(*vmat2:GFP*) zebrafish contained a reporter gene for GFP under the expression of the vesicular monoamine transporter (*vmat2* also known as *slc18a2*) and were used to visualize 5-HT-containing NECs. 5HT_3_ receptors in the gill were identified using polyclonal rabbit anti-5HT_3B_ receptor at 1:100 (RRID: AB_11105287 cat. no. BS-4289R, Bioss Inc., Woburn, MA, USA) and visualized using 1:100 Alexa 594 goat anti-rabbit secondary antibody.

Whole-mount preparations were examined using an upright microscope platform (H101A ProScan, Olympus, Canada) with motorized XYZ control and a confocal scanning system (FV100 BX61 LSM, Olympus) equipped with continuous laser lines at 405 nm, 488 nm and 561 nm.

### Acclimation to chronic hypoxia

Adult zebrafish were placed in 2.5-liter tanks and acclimated to hypoxia (35 mmHg) for 48 h. A group of control zebrafish were simultaneously maintained in an identical tank at normoxia (∼160 mmHg) for the same period of time. For acclimated fish, water PO_2_ was gradually lowered over 8 h at a rate of ∼16 mmHg h^-1^ by introducing a mixture of compressed air and 100% N_2_ from a gas mixer (Pegas 4000 MF; Columbus Instruments, Columbus, OH, USA) and delivered through a porous air stone. Water temperature was kept at 28°C by placing tanks in a temperature-controlled water bath and 50% water changes were performed in both tanks every day using hypoxic water for hypoxia treated groups and normoxic water for control groups.

### Quantitative polymerase chain reaction

Whole gill baskets were removed from zebrafish acclimated to hypoxia or normoxic controls after 48 h and immediately frozen on dry ice. Total RNA from gill baskets was extracted using Trizol reagent (Life Technologies, Carlsbad, CA, USA) and quantified using a NanoDrop 2000c UV–Vis Spectrophotometer (Thermo Fisher Scientific, Waltham, MA, USA). cDNA was generated using 1000 ng of RNA from whole gill baskets using QuantiTect Reverse Transcription Kit (Qiagen, Toronto, ON, Canada) following manufacturer protocols. To check for genomic DNA contamination, a no-template negative control and a no reverse transcriptase negative control were included. mRNA gene expression of nAChR subunits α2, α3, α4, α6, α7, β2, β3, and β4 (*chrna2a*, *chrna3*, *chrna4*, *chrna6*, *chrna7*, *chrnb2a*, *chrnb3*, and *chrnb4*) and serotonin receptor 5HT_3_ (*htr3a* and *htr3b*) in the gill was assessed by real-time reverse transcription quantitative polymerase chain reaction (RT-qPCR) using SsoAdvanced Universal SYBR Green Supermix (Bio-Rad, Mississauga, ON, Canada). Expression of reference gene, elongation factor 1a (*ef1a*), was stable between gill baskets of normoxia and hypoxia groups and was therefore used to normalize mRNA expression of all genes (Reed et al., 2024). Standard curves were generated using a serial dilution of pooled cDNA to optimize primer reaction conditions. Quantitative polymerase chain reaction included 1 μl cDNA, 1 μl specific forward primer, 1 μl specific reverse primer, 7 μl nuclease-free water, and 10 μl Universal SYBR Green Supermix for a total reaction volume of 20 μl. Each reaction included an initial step at 98°C to activate the enzymes in the mix, followed by 40 repeats of a denaturation step at 95°C and an annealing/extension step at the optimized temperature for each primer pair (Tables 1 and 2). All primers were designed using Primer3 software (Untergasser et al., 2012), and reaction efficiencies of 90-110%, and R2 values >0.98, were used as quality control. A melt step from 66 to 95°C in 0.5°C increments was included at the end of the reaction to check that a single amplicon was produced. Each biological sample was run in triplicate, and the relative abundance of mRNA was calculated using the ΔΔCt method (Livak and Schmittgen, 2001). Relative transcript abundances are expressed as fold-change compared to normoxic control groups.

**Table 1.**
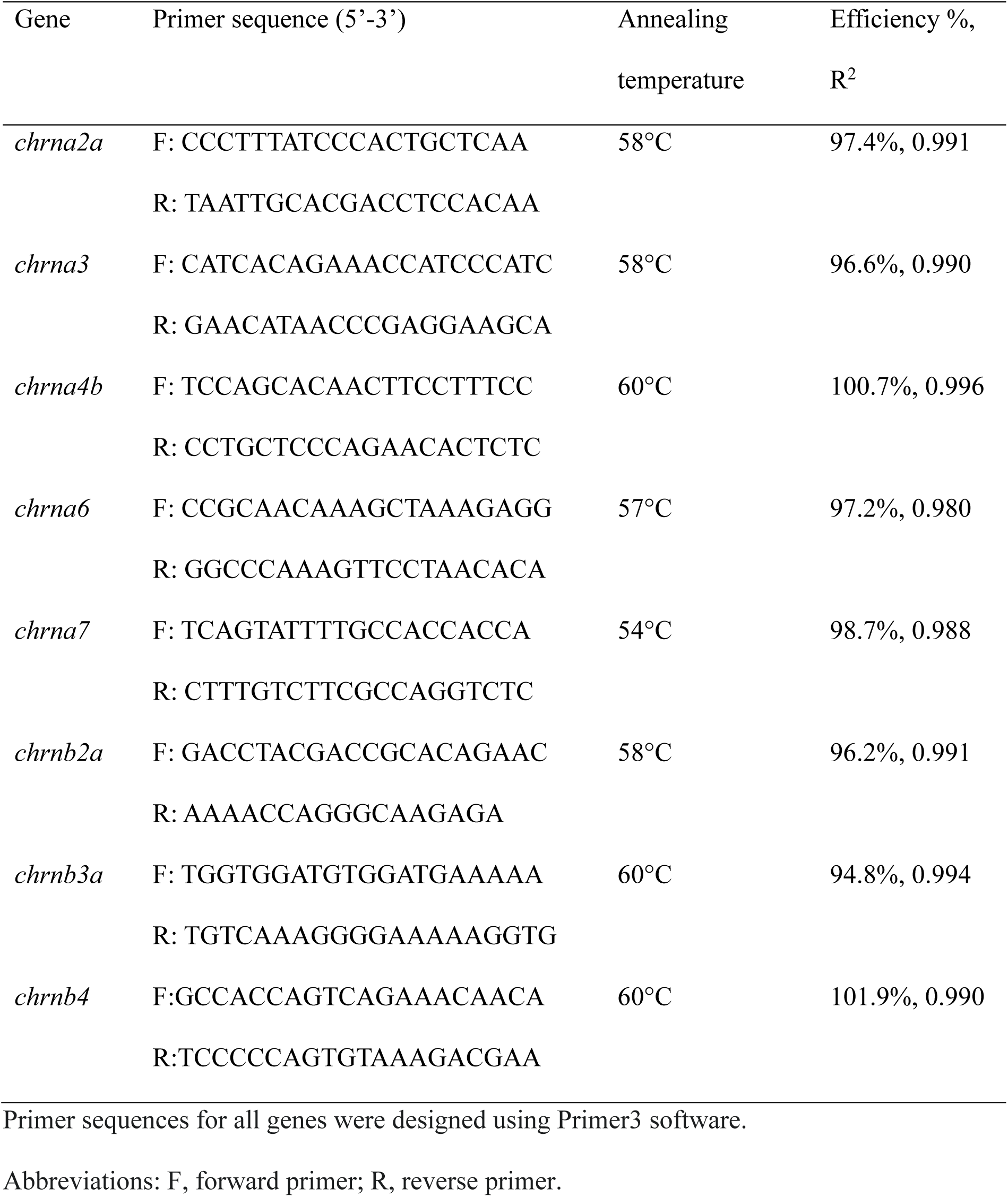
Nicotinic receptor subunit primer pair conditions used for mRNA quantification by real-time reverse transcription polymerase chain reaction.

**Table 2.**
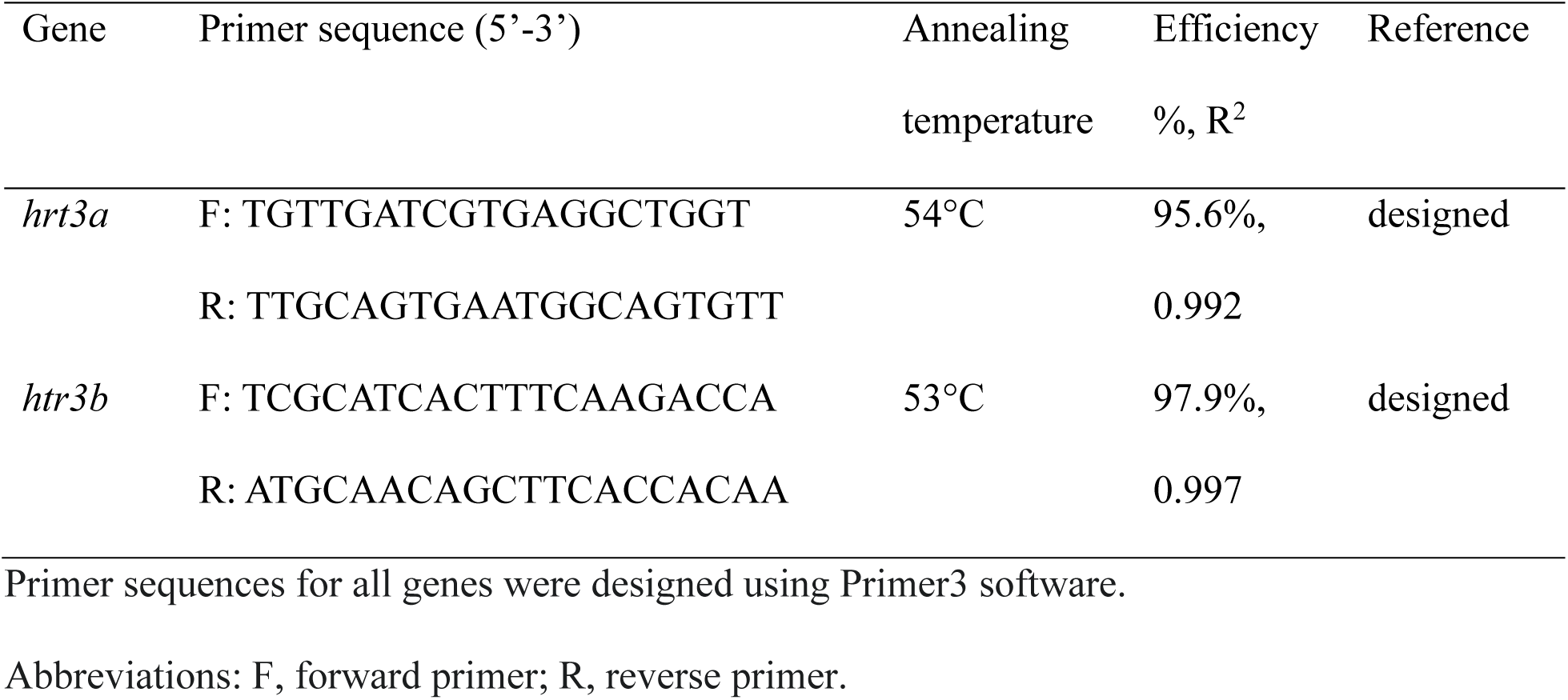
Serotonin receptor primer pair conditions used for mRNA quantification by real-time reverse transcription polymerase chain reaction.

### Statistical analysis

For all reported data, sample size (n) refers to individual cells. While multiple gill arches were assessed per animal, only one cell from each arch was included in the analysis to avoid repeated exposures or treatments in the same tissue. For each recording, the baseline fluorescence was calculated as the average fluorescence intensity of a cell for the first 30 s in normoxia. All fluorescence values were divided by the baseline to evaluate changes in fluorescence intensity over time throughout a single recording. Statistical analysis for gill and whole animal Ca^2+^-imaging recordings were carried out using the Wilcoxon matched-pairs signed rank test for paired comparisons or the Mann-Whitney U test for unpaired comparisons with Prism v9.5.1 (GraphPad Software Inc., San Diego, CA, USA). For comparison of paired samples exposed to three or more treatments a Friedman test with Dunn’s multiple comparison test was used. Dose-responses were normalized to the maximum concentration of each drug. For estimation of EC_50_, a line constrained from the origin at zero was fit to the data using a nonlinear [agonist/antagonist] vs. normalized response model with least squares following the equation: y=100*(X^HillSlope^)/(EC50^HillSlope^ + X^HillSlope^) using Prism v9.5.1). All data were expressed as means ± standard deviation (SD). Statistical analysis for all qPCR expressional data was carried out using the Mann–Whitney *U* test with Prism v9.5.1.

## Results

### Cholinergic activation increased intracellular Ca^2+^ concentration ([Ca^2+^]_i_) in chain neurons

To evaluate a role for ACh as an excitatory neurotransmitter in the ChN response to hypoxia, we first examined changes in ChN [Ca^2+^]_i_ in response to drugs that target cholinergic receptors. In whole-gill recording, exogenous addition of ACh, at concentrations between 50 and 500 μM, caused a dose-dependent increase in [Ca^2+^]_i_ in ChNs (Fig. 1A,B) with an EC_50_ of 168.3 µM (Fig. 1C). In a similar manner, the ACh receptor agonist, nicotine, caused a dose-dependent increase in [Ca^2+^]_i_ (Fig 1D,E) with an EC_50_ of 37.68 µM (Fig. 1F). Further, blockade of nicotinic ACh receptors using hexamethonium reduced the ChN response to hypoxia in a dose-dependent manner (Fig. 2A,B). In experiments where the ChN was first exposed to hypoxia alone, during a second bout of hypoxia the [Ca^2+^]_i_ response was reduced by 31.25% with 25 µM (Wilcoxon matched-pairs signed rank test, p=0.0623, n=5), by 46.91% with 50 µM (p=0.0313, n=6), by 51.16% with 100 µM (p=0.0156, n=6), by 68.89% with 200 µM (p=0.0156, n=6), and by 74.66% with 300 µM hexamethonium (p=0.0313, n=5). The hypoxic response, with the addition of 200 and 300 µM hexamethonium, was not significantly different from baseline fluorescence in normoxia (P>0.9999, n=5-6), indicating the hypoxia-induced increase in [Ca^2+^]_i_ was completely abolished by hexamethonium at these concentrations. The EC_50_ of hexamethonium was 79.74 µM (Fig. 2C).

**Figure 1.**
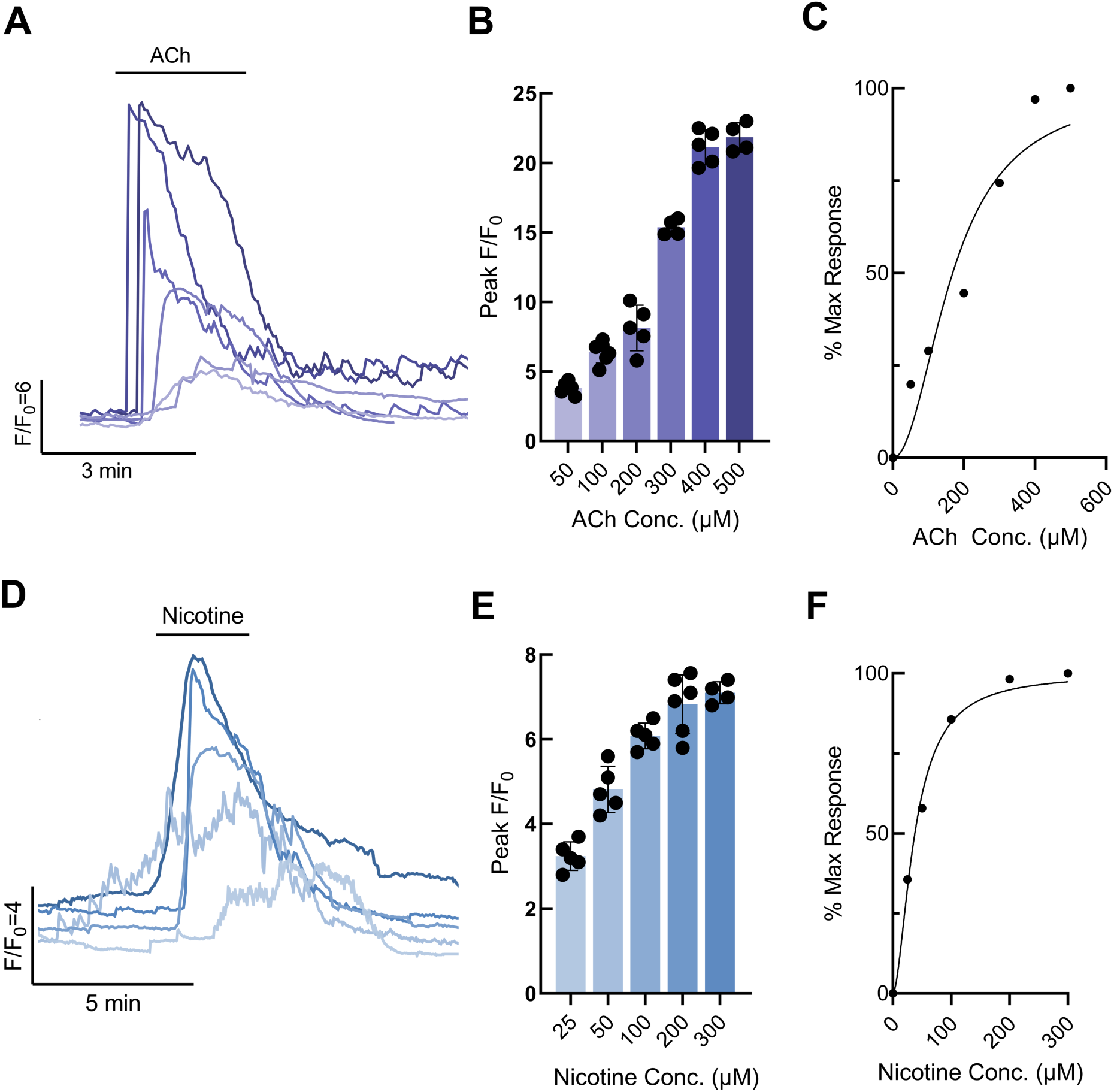
Postsynaptic neurons displayed a dose-dependent increase in intracellular Ca^2+^ concentration ([Ca^2+^]_i_) in response to exogenous acetylcholine and nicotine. (A) Overlayed Ca^2+^ imaging traces from 6 chain neurons (ChNs) exposed to 50-300 μM acetylcholine (ACh). Scale indicates time (min) and relative changes in fluorescence (F/F_0_) corresponding to changes in [Ca^2+^]_i_. (B) Summary data with mean ± S.D. from ChNs as treated in (A) (n=4-6). (C) Dose-response from summary data in (B) showing concentration versus percent maximum response. Data points were fit using a nonlinear [agonist] vs. normalized response model with least squares following the equation: y=100*(X^HillSlope^)/(EC50^HillSlope^ + X^HillSlope^). The effective half maximal concentration (EC_50_) for ACh was 168.30 μM. (D) Overlayed Ca^2+^ imaging traces from 5 ChNs exposed to 25-300 μM nicotine. Scale indicates time (min) and relative changes in fluorescence (F/F_0_) corresponding to changes in [Ca^2+^]_i_. (E) Summary data with mean ± S.D. from ChNs as treated in (D) (n=4-6). (F) Dose-response from summary data in (E) with linear fit described for (C) showing concentration versus percent maximum response. EC_50_ for nicotine was 37.68 μM.

**Figure 2.**
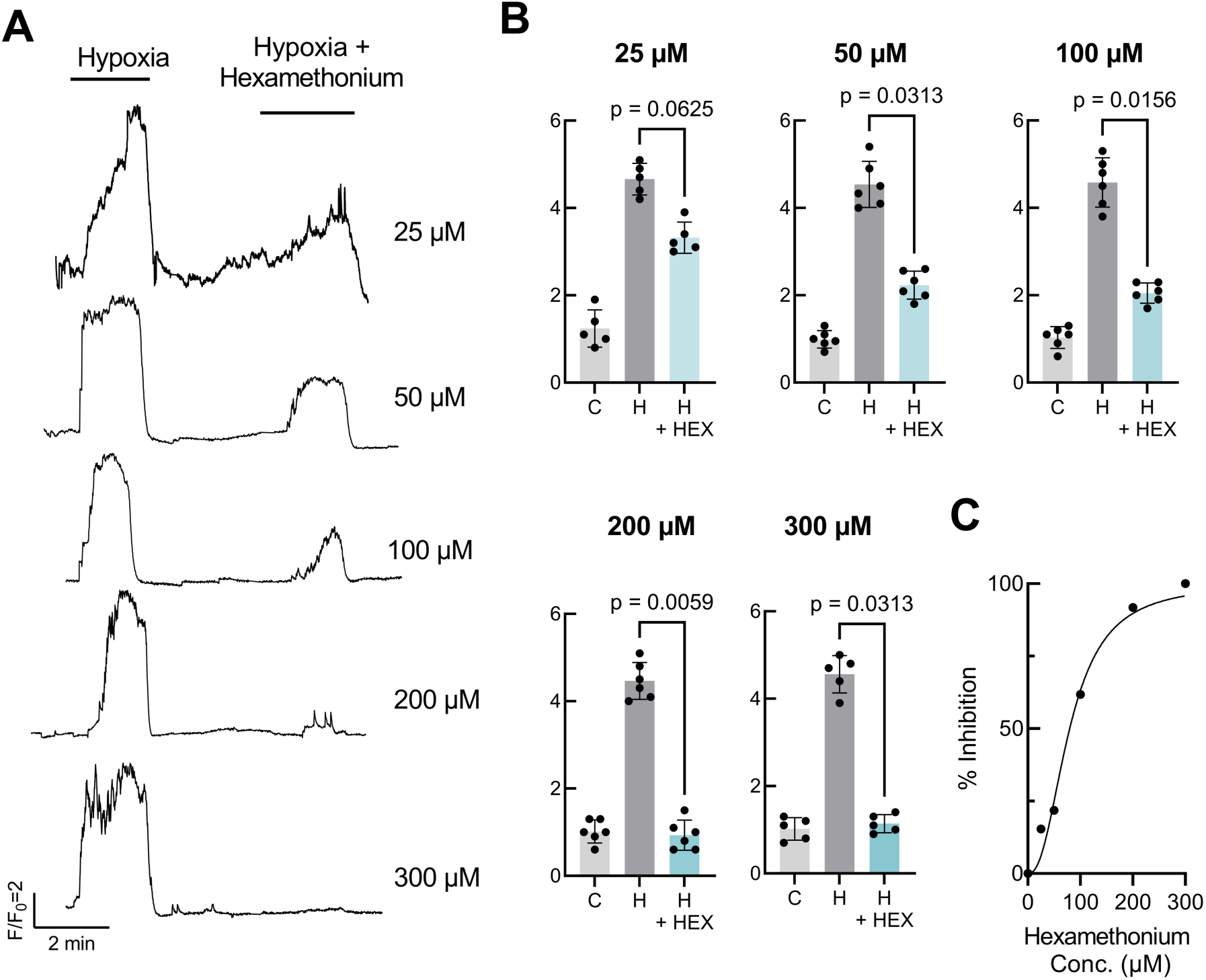
**The postsynaptic neuron hypoxia-induced increase in intracellular calcium concentration ([Ca^2+^]_i_) was abolished by the nicotinic acetylcholine receptor antagonist, hexamethonium**. (A) Ca^2+^ imaging traces from 5 chain neurons (ChNs) each exposed to a 2-min bout of hypoxia followed by recovery in normoxia and a second bout of hypoxia in the presence of increasing concentrations of hexamethonium (25-300 μM). (B) Summary data with mean ± S.D. from ChNs as treated in (A) comparing relative changes in fluorescence (F/F_0_) during the following conditions: baseline control fluorescence in normoxia (C), first hypoxia exposure (H), and second hypoxia exposure with hexamethonium (H+HEX). The hypoxic response (increase in [Ca^2+^]_i_) was reduced by 31.25% with 25 μM hexamethonium (Wilcoxon matched-pairs signed rank test, p=0.0623, n=5), by 46.91% with 50 μM hexamethonium (p=0.0313, n=6) by 51.16% with 100 uM hexamethonium (p=0.0156, n=6), by 68.89% with 200 μM hexamethonium (p=0.0156, n=6), and by 74.66% with 300 μM hexamethonium (p=0.0313, n=5). The hypoxic response with the addition of 200 or 300 μM hexamethonium was not significantly different than baseline activity in normoxia (P>0.9999, n=6). (C) Dose response generated from summary data in (B) showing concentration versus percent maximum response. Data points were fit using a nonlinear [antagonist] vs. normalized response model with least squares following the equation: y=100*(X^HillSlope^)/(EC50^HillSlope^ + X^HillSlope^). The EC_50_ for hexamethonium was 79.74 μM.

### Acetylcholine and nicotinic ACh receptors were localized in the gills

The experiments presented in Figure 2 suggested that ACh was endogenously released from chemoreceptors during hypoxia and that postsynaptic receptors on ChNs mediate cholinergic responses. Using immunohistochemistry and confocal imaging, we localized Ach immunoreactivity (IR). ACh-IR colocalized with SV2 (Fig. 3A-C), a common marker for neurosecretory cells in the gills, including NECs that do not contain 5-HT (Jonz and Nurse, 2003). Moreover, ACh-IR was found in a number of GCaMP-positive NECs (Fig. 3D-F), which were previously shown to be oxygen-sensitive (Reed and Jonz, 2025). ACh-IR did not co-localize with expression of VMAT2 (Fig. 3G-I)—an established marker of serotonergic NECs (Pan et al., 2021)—suggesting that ACh-IR cells were an independent population of NEC that did not contain 5-HT and were most abundant in the tips of filaments (Fig. 3J-L). These observations led us to propose the presence of two distinct populations of NECs. We refer to these as: S-type NECs, containing serotonin (5-HT) and SV2-positive synaptic vesicles; and A-type NECs, with ACh and SV2-positive vesicles.

**Figure 3.**
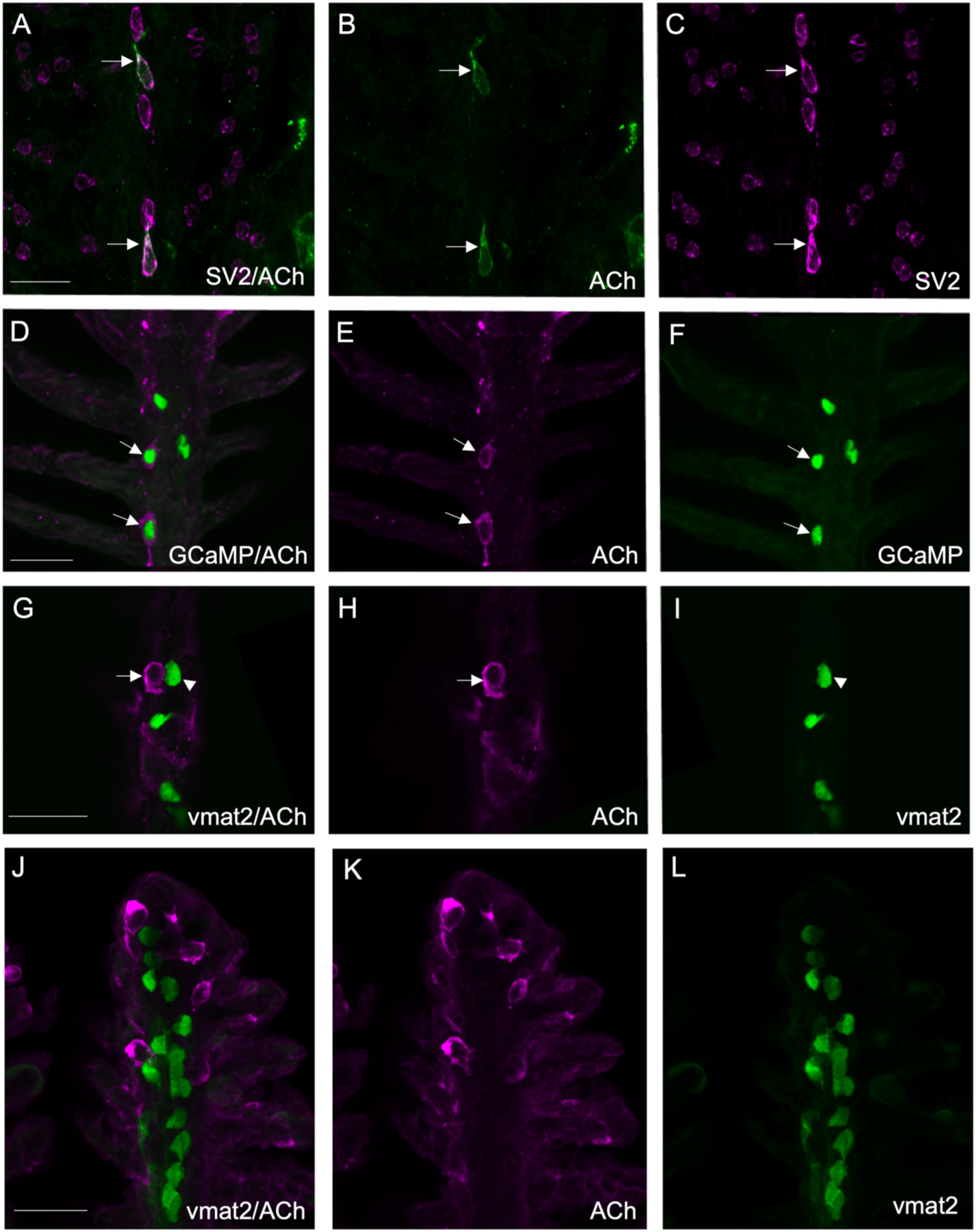
Confocal imaging of immunohistochemical localization of acetylcholine-positive neurosecretory cells in the gill filament epithelium. (A-C) Co-labeling of acetylcholine (ACh, green) in neurosecretory cells containing synaptic vesicle protein-2 (SV2, magenta, arrows). (B,C) ACh and SV2 labeling shown separately. Cells labeled with both markers are indicated by arrows. Scale bar in A=25 µM and applies to B and C. (D-F) Confocal imaging of immunohistochemical localization of ACh (magenta) with GCaMP-positive neuroepithelial cells (NECs, green, arrows) in gill filaments of Tg(*elavl3*:GCaMP6s) zebrafish. (E,F) ACh and GCaMP labeling shown separately. Cells labeled with both markers are indicated by arrows. Scale bar in D=25 µM and applies to E and F. (G-I) Co-labeling of ACh (magenta) and the vesicular monoamine transporter-2 (*vmat2*, green). Scale bar in G=20 µM and applies to H and I. ACh-positive neurosecretory cells (arrow) did not co-localize with *vmat2*-positive NECs (arrowhead). (H,I) ACh and *vmat2* labeling shown separately. (J-L) Co-labeling of ACh (magenta) and the vesicular monoamine transporter-2 (*vmat2*, green). Scale bar in J=25 µM and applies to K and L. *vmat2*-positive NECs were localized along the midline of the filament, whereas ACh-positive neurosecretory cells lied away from the midline and were most concentrated at the tips of the filaments. (K,L) ACh and *vmat2* labeling shown separately.

Using Ca^2+^ imaging in isolated gills, we demonstrate that hypoxia enhanced the ChN response to low concentrations of nicotine at 25 µM (Wilcoxon matched-pairs signed rank test, p=0.0313, n=5) and 50 µM (p=0.313, n=5), but at higher concentrations of nicotine (100-300 µM) the ChN response was not significantly enhanced by hypoxia (n=4-5, Fig. 4A,B). The amount of fluorescence signal added by hypoxia drastically declined at concentrations of nicotine above 100 µM (Fig. 4C). This occlusion of the enhancing effect of hypoxia at higher nicotine concentrations suggests the endogenous excitatory neurotransmitter released by A-type NECs and stimulating ChNs was acting through nAChRs.

**Figure 4.**
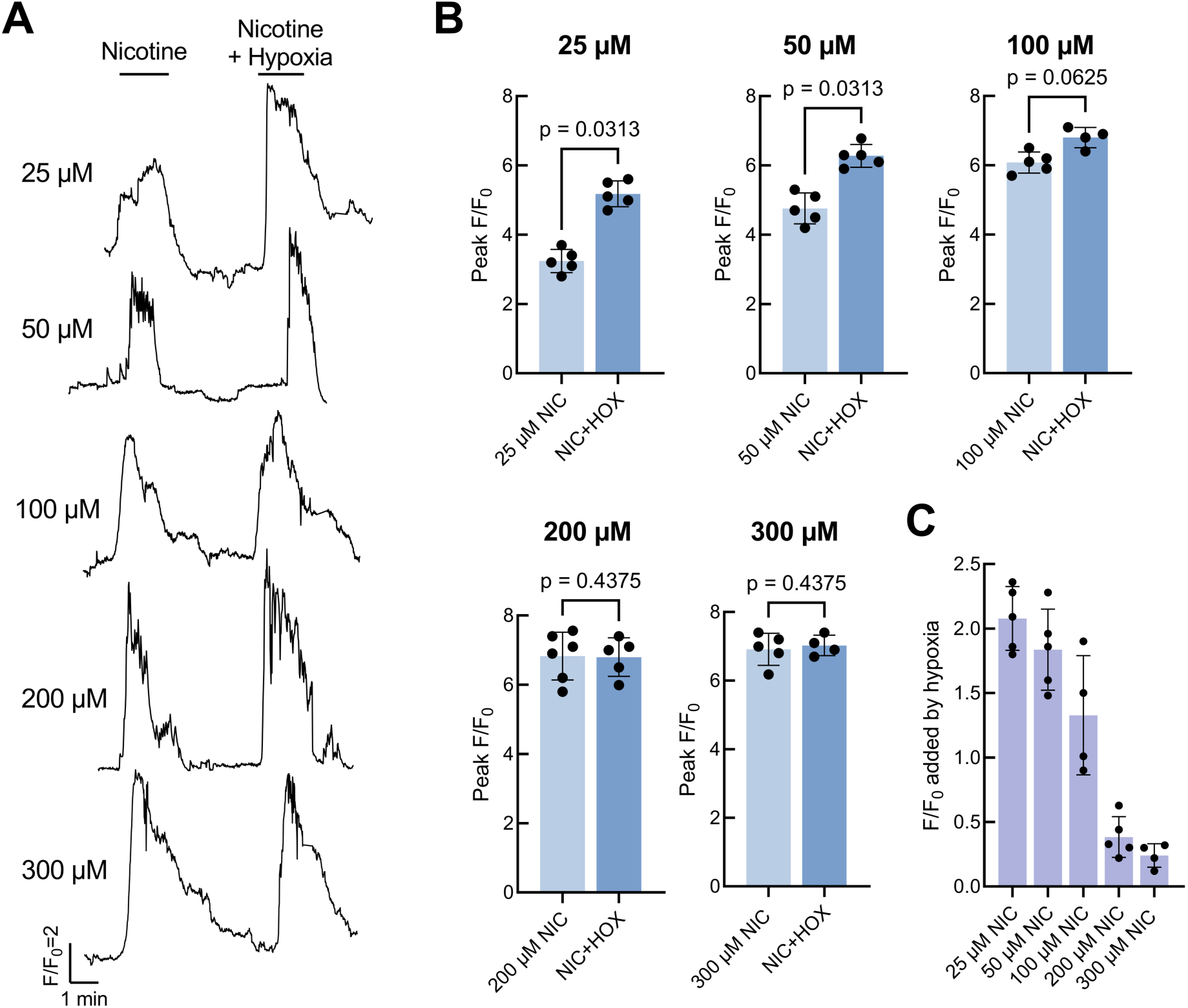
The additive effect of hypoxia on the neuronal increase in intracellular calcium concentration ([Ca^2+^]_i_) in response to nicotine is abolished at high nicotine concentrations. (A) Ca^2+^imaging traces from 5 GCaMP-containing chain neurons (ChNs) comparing the [Ca^2+^]_i_ response to increasing concentrations of nicotine with and without the addition of hypoxia. The same concentration of nicotine (indicated at left) was applied twice in each trace. (B) Summary data with mean ± S.D. from ChNs as treated in (A) comparing the effect of hypoxia on each concentration of nicotine. Hypoxia had a significant enhancing effect on the ChN response to 25 μM (Wilcoxon matched-pairs signed rank test, p=0.0313, n=5) and 50 μM nicotine (p=0.313, n=5) but did not significantly enhance 100-300 μM nicotine (n=4-5). (C) Summary data with mean ± S.D. from (B) showing the F/F_0_ corresponding to changes in [Ca^2+^]_i_ added by hypoxia at each concentration of nicotine. The additive effect of hypoxia is occluded at high nicotine concentrations.

In qPCR experiments using isolated gill tissue, we confirmed expression of genes *chrna2a, chrna3, chrna4, chrna6, chrna7, chrb2a*, *chrnb3* and *chrnb4*, encoding nicotinic receptor subunits α2, α3, α4, α6, α7, β2, β3 and β4, respectively (Fig. 5). 48 h of hypoxia exposure upregulated expression of many of these genes, including *chrna2a* (Mann-Whitney test, p=0.0006, n=7), *chrna3* (p=0.0012, n=7), *chrnb2a* (p=0.0262, n=7) and *chrnb4* (p=0.0006, n=7). Importantly, these data showed that the gene encoding the α2 subunit displayed the greatest relative increase in expression following hypoxia, suggesting a possible role in hypoxia signaling.

**Figure 5.**
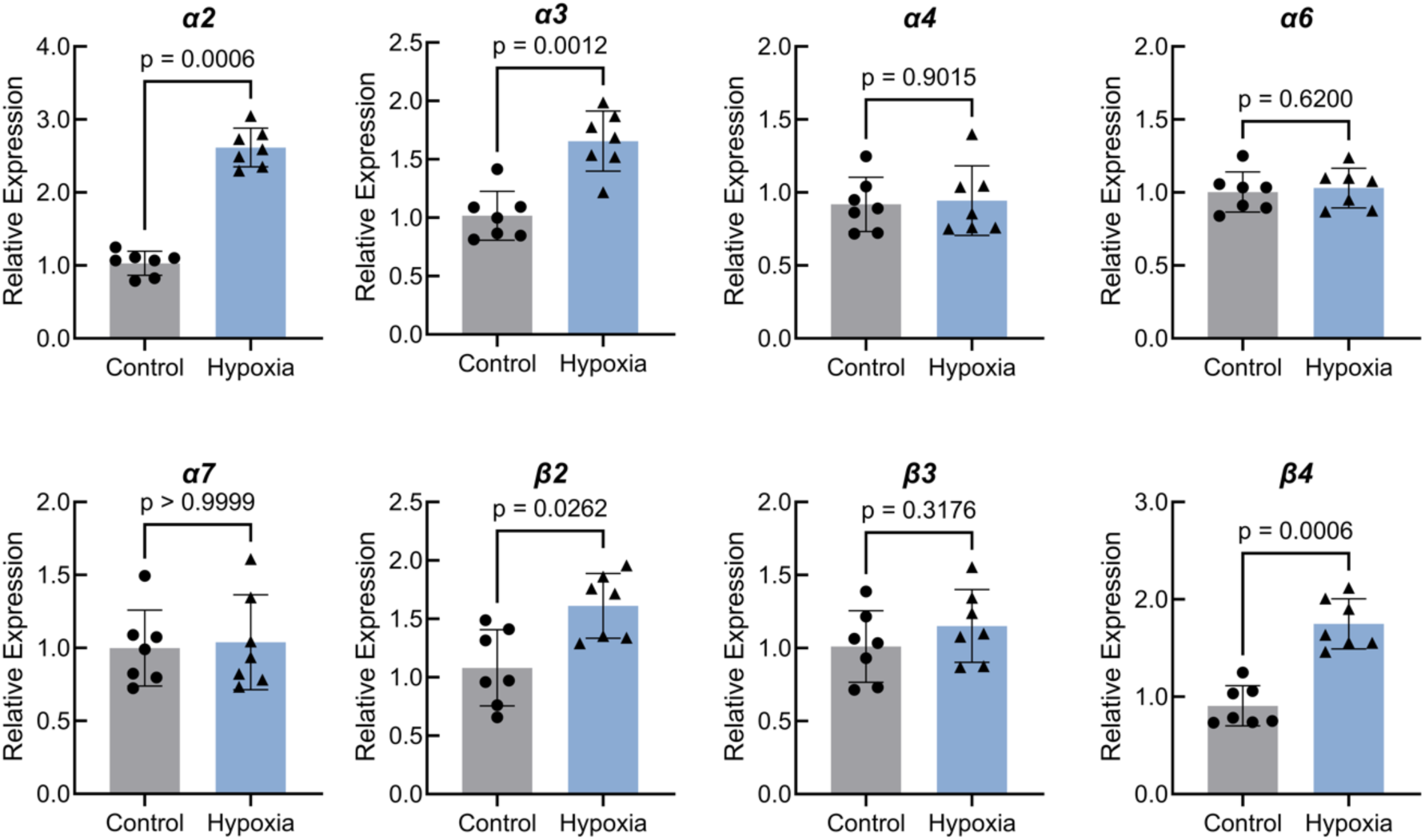
Relative mRNA expression of genes encoding nicotinic acetylcholine receptor (nAChR) subunits in isolated gill tissue. Expression of *chrna2a, chrna3, chrb2a*, and *chrnb4* (encoding nAChR subunits α2, α3, β2, and β4) increased following 48 h of chronic hypoxia exposure. Data were normalized to the mRNA abundance of the reference gene, *ef1a*. Mean ± S.D. is shown and data were analyzed using a Mann–Whitney *U* test (n = 7).

Using confocal microscopy, we localized nAChR α2 subunit-IR to nerve fibers travelling along the midline of the gill filament. α2 subunit labeling was punctuated with regions of increased fluorescence intensity and colocalized with anti-zn-12 (Fig. 6A-C, arrows), a zebrafish neuronal marker that also labels other nerve fibers and ChNs. Rotation by 90° showed that nerve fibers projecting from ChNs coursed around the filament artery to the superficial layer, where α2 subunit-IR was found (Fig. 6D-F, arrowhead). When AChR α2 subunit immunoreactivity was combined with anti-SV2, it was evident that α2 subunit-IR puncta occurred along the nerve fibers in close proximity with SV2-positive NECs (Fig. 6G-I).

**Figure 6.**
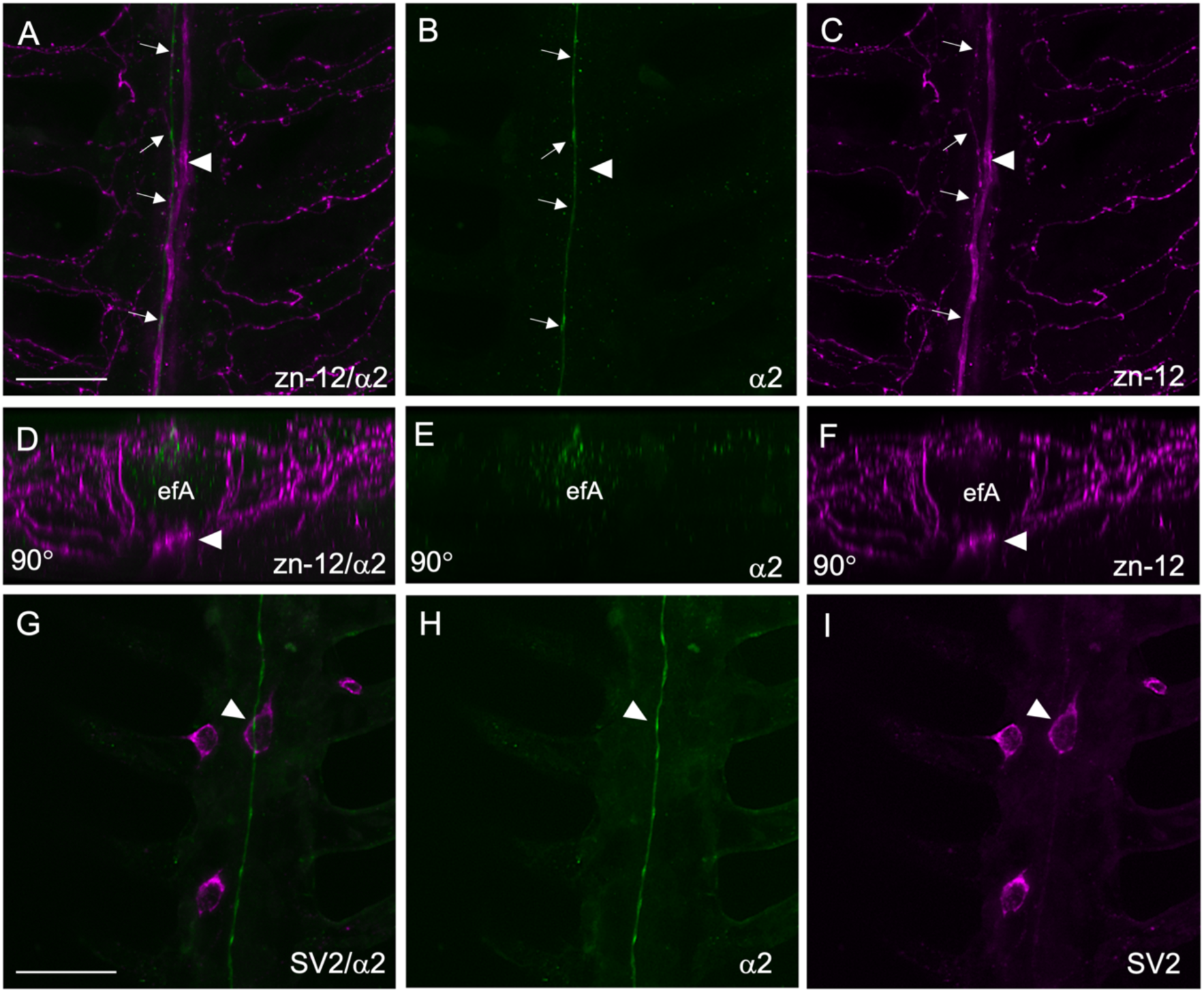
Nerve fibers containing acetylcholine receptor (AChR) subunit α2 arise from chain neurons and contact neuroepithelial cells (NECs). (A) Immunolabeling of AChR α2 subunit (green, arrows) was punctuated with regions of increased fluorescence intensity and colocalized with nerve fibers labeled with anti-zn-12 (magenta). Scale bar = 20 μm and applies to panels B-F. (B,C) AChR α2 subunit and zn-12 labeling shown separately. (D-F) Images from A-C tilted 90° to show spatial separation between the α2-positive nerves, which lie superficial and lateral to the efferent filament artery (efA) and the deeper chain neurons (ChNs, arrowhead). (G) SV2 labeled a NEC (magenta) in the gill epithelium in close association with AChR subunit α2-positive nerve fiber (green, arrowhead). Scale bar = 25 μm and applies to H and I. (H,I) α2 subunit and SV2 labeling shown separately.

### Cholinergic stimulation in the gills led to time-dependent activation of vagal sensory ganglia

To determine whether the hypoxic signal generated in the gill was transmitted towards the central nervous system, we developed a preparation to investigate ganglionic activity in the head of intact Tg(*elavl3*:GCaMP6s) larvae (Fig. 7A-C). Each gill arch receives sensory autonomic innervation from an epibranchial ganglion, which projects to the larger nodose ganglion of the vagus nerve. Both ganglia are collectively referred to as the vagal sensory ganglia. Exposure of intact larvae to hypoxia corresponded with a significant rise in [Ca^2+^]_i_ from all four epibranchial ganglia (Fig.7D), consistent with increased activity in all gill arches. Similarly, exposure of intact larvae to hypoxia corresponded with a significant rise in [Ca^2+^]_i_ from the nodose ganglion (Fig. 7E, Wilcoxon matched-pairs signed rank test, p=0.156, n=7).

**Figure 7.**
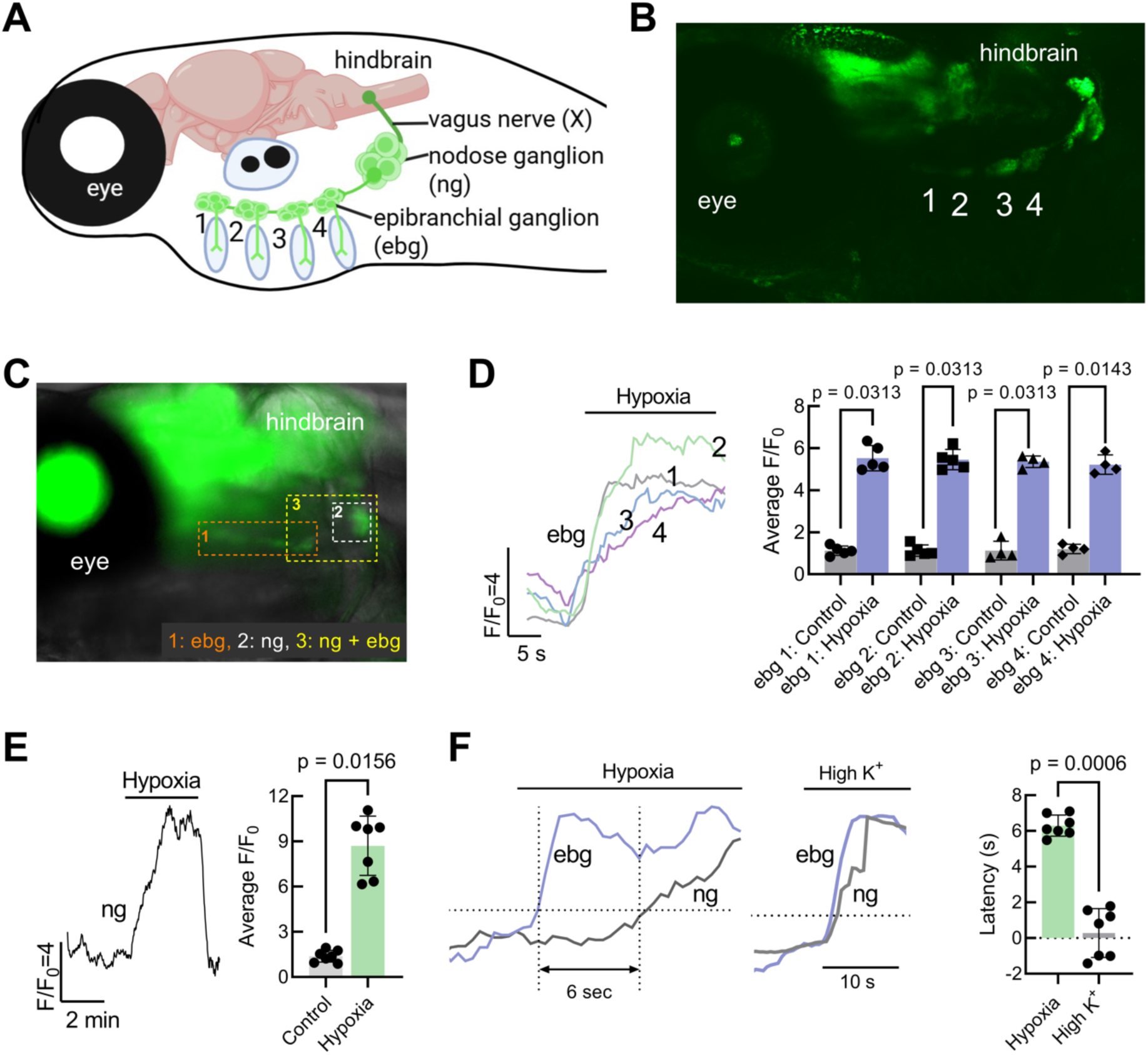
Larval preparation for evaluating changes in intracellular calcium concentration ([Ca^2+^]_i_) associated with hypoxia in vagal sensory ganglia. (A) Schematic of vagus nerve (X) innervation of the gills, including the large nodose ganglion (ng) and four epibranchial ganglia (ebg, 1-4) above each gill arch in a larval zebrafish. Created using Biorender. (B) Confocal image from a 15 days post-fertilization larval Tg(*elavl3*:GCaMP6s) zebrafish expressing GFP (green) associated with GCaMP in the nodose ganglion and epibranchial ganglia (1-4). (C) Overlay of brightfield and green fluorescence (488 nm) image of the vagal ganglia *in vivo* containing GCaMP from a larval Tg(*elavl3*:GCaMP6s) zebrafish. Dashed boxes indicate areas of focus for subsequent Ca^2+^ imaging experiments. Recording region 1 (orange dashed box) included all four epibranchial ganglia (ebg) to be observed simultaneously. Recording region 2 (white dashed box) included only the nodose ganglion (ng). Recording region 3 (yellow dashed box) included the nodose ganglion and the fourth epibranchial ganglion (ng + ebg) to be observed simultaneously. (D, left) Simultaneous Ca^2+^ imaging recording from all four epibranchial ganglia (ebg) during hypoxia exposure. (D, right) Summary data with mean ± S.D. for hypoxia-induced [Ca^2+^]_i_ increases in epibranchial ganglia (Wilcoxon matched-pairs signed rank test, n=4-5). (E) Ca^2+^ imaging trace from the nodose ganglion (ng) during whole-animal exposure to hypoxia (left). Summary data (mean ± SD) for hypoxia-induced [Ca^2+^]_i_ increases in larval nodose ganglion (right, Wilcoxon matched-pairs signed rank test, p=0.0156, n=7). (F, left) Simultaneous Ca^2+^ imaging trace from an epibranchial ganglion (ebg, blue) and nodose (ng, grey) in response to hypoxia. (F, middle) Simultaneous Ca^2+^ imaging trace from an epibranchial ganglion (ebg, blue) and nodose (ng, grey) in response to high K^+^ concentration. (F, right) Summary (mean ± SD) of the time delay between the onset of epibranchial and nodose [Ca^2+^]_i_ responses to hypoxia and high K^+^ in 7 fish. Onset of the [Ca^2+^]_i_ response was characterized as the time where relative fluorescence (F/F_0_) first doubled. Hypoxia exposure resulted in a larger time delay between activation of the epibranchial and nodose ganglia compared to high K^+^ (Mann-Whitney test, p=0.0006, n=7).

Next, we simultaneously evaluated [Ca^2+^]_i_ responses from the posterior epibranchial ganglion above the fourth gill arch and the nodose ganglion. When whole larvae were exposed to hypoxia, both ganglia displayed an increase in [Ca^2+^]_i_ (Fig. 7F). Notably, when recorded simultaneously, we observed that the epibranchial ganglion, closest to the gill, was always first to respond to hypoxia, followed by the nodose ganglion. When measuring the onset of the [Ca^2+^]_i_ response as the time where F/F_0_ doubled, the response in the nodose ganglion occurred after a latency of 6.3 ± 0.6 s (n=7) compared to the epibranchial ganglion (Fig. 7F). In contrast, both populations of ganglionic neurons were activated within 0.28 ± 1.37 s (n=7) of each other, and in no specific order, in response to extracellular solution containing high K^+^ to directly induce membrane depolarization in both neuronal populations. This served as a control experiment to demonstrate that when both ganglia responded directly to the same stimulus (i.e. high K^+^), a sequence of responses could not be determined. A significant difference between the latency in activation of the epibranchial versus nodose ganglia during hypoxia was found when compared to high K^+^ (Mann-Whitney test, p=0.0006, n=7). This indicated that hypoxia signaling involves directional transmission from the gill towards the nodose, and that vagal sensory ganglia were unlikely to respond directly to hypoxia.

Exogenous addition of ACh or nicotine, which activated ChNs in the gill in a manner similar to that of hypoxia (Figs. 1,4), evoked an increase in [Ca^2+^]_i_ from the nodose ganglion in recordings from intact larvae (Fig. 8A, Wilcoxon matched-pairs signed rank test, p=0.0156, n=7 and Fig. 8B, p=0.0313, n=7). Further, a significant difference was found in the latency between ACh-induced responses of the epibranchial and nodose ganglia (8.1 ± 0.7 s) compared to the latency between ganglionic responses to direct activation by high K^+^ (0.28 ± 1.37 s; Fig. 8C, Mann-Whitney test, p=0.0006, n=7). Likewise, we observed a significant difference in the latency between ganglionic responses after administration of nicotine (8.8 ± 0.9 s) or high K^+^ (0.28 ± 1.37 s; Fig. 8D, Mann-Whitney test, p=0.0006, n=7). These results suggest that, like hypoxia, ACh and nicotine activated ChNs in the gill, and this signal was transmitted to the vagal sensory ganglia.

**Figure 8.**
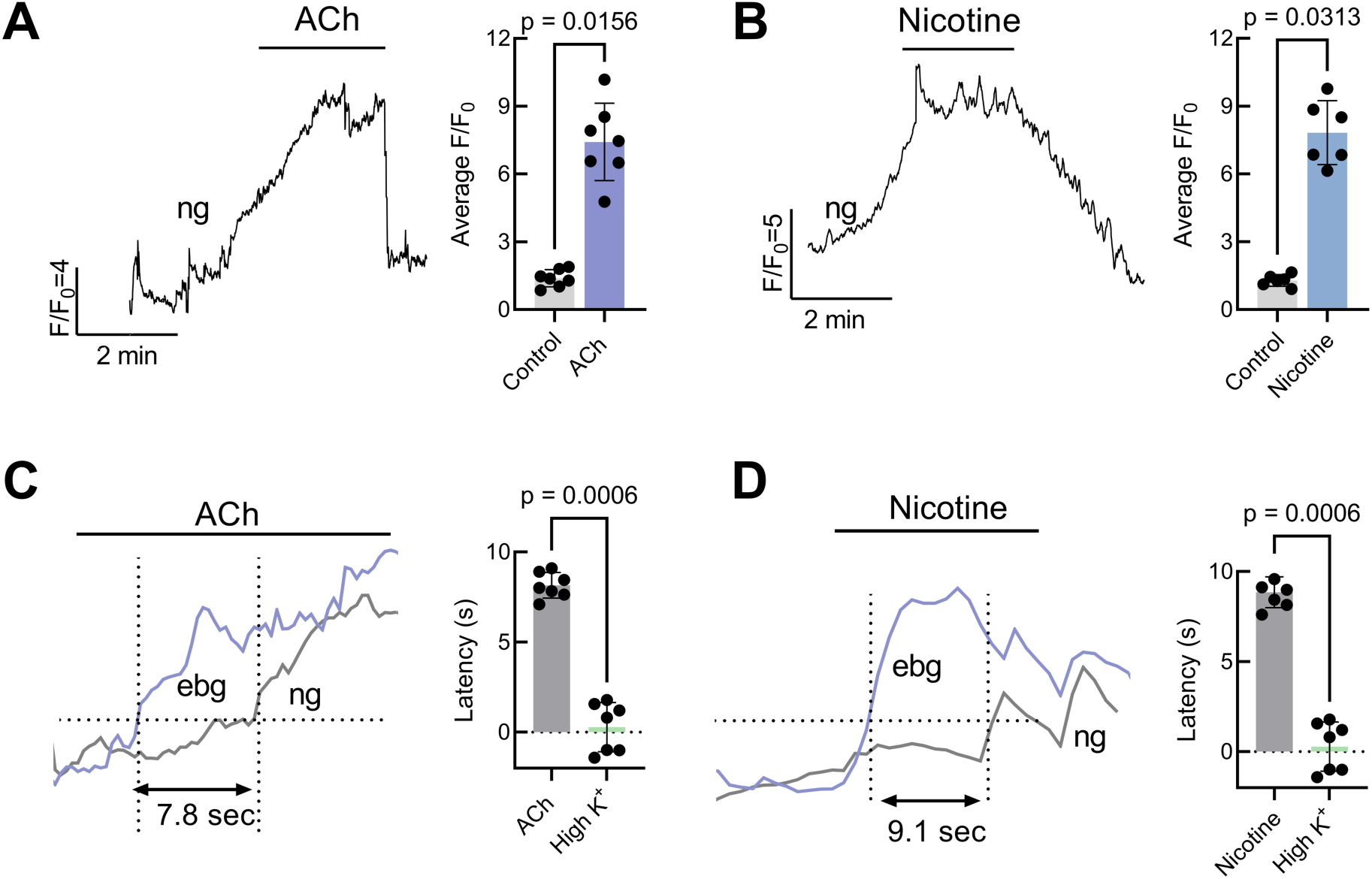
Vagal sensory ganglia increased intracellular calcium concentration ([Ca^2+^]_i_) in response to acetylcholine and nicotine in whole-animal larval Ca^2+^ imaging recordings. (A) Ca^2+^ imaging trace from a larval GCaMP-containing nodose ganglion (ng) during exposure to 50 μM ACh (left). Scale indicates time (min) and relative changes in fluorescence (F/F_0_) corresponding to changes in [Ca^2+^]_i_. At right, mean ± SD F/F_0_ nodose ganglion responses to ACh (Wilcoxon matched-pairs signed rank test, p=0.0156, n=7). (B) Ca^2+^ imaging trace from a larval GCaMP-containing nodose ganglion (ng) during exposure to 50 μM nicotine (left). Scale indicates time (min) and relative changes in fluorescence (F/F_0_) corresponding to changes in [Ca^2+^]_i_. At right, mean ± SD F/F_0_ nodose responses to nicotine (Wilcoxon matched-pairs signed rank test, p=0.0313, n=7). (C) Simultaneous Ca^2+^ imaging recording showing onset time for the epibranchial (ebg) and nodose (ng) ganglia in response to 50 μM ACh (left). Onset of the [Ca^2+^]_i_ response was characterized as the time where relative fluorescence first doubled (F/F_0_=2). At right, summary data showing mean ± SD time between onset of epibranchial and nodose ganglia [Ca^2+^]_i_ responses to ACh compared to high K^+^ control (Mann-Whitney test, p=0.0006 n=7). (D) Simultaneous Ca^2+^ imaging recording showing response onset time for the epibranchial (ebg) and nodose (ng) ganglia in response to 50 μM nicotine (left). Onset of the [Ca^2+^]_i_ response was characterized as the time where relative fluorescence (F/F_0_) first doubled. At right, summary data showing mean ± SD time between epibranchial and nodose ganglia [Ca^2+^]_i_ responses to nicotine compared to high K^+^ control (Mann-Whitney test, p=0.0006, n=7). Both ACh and nicotine produced a longer delay between activation of epibranchial and nodose ganglia compared to the high K^+^ control.

### Serotonergic hypoxic signaling in the gill occurred independently of the chain neuron pathway

Given the prevalence of 5-HT in the gill, and that gill neurons in zebrafish display strong expression of 5-HT_3_ receptors (Pan et al., 2022), we investigated the potential effects of 5-HT on hypoxic signaling in both our whole-animal larval preparation and isolated gill preparation. Both 5-HT and the 5-HT_3_ receptor agonist, phenylbiguanide, produced an increase in nodose ganglion [Ca^2+^]_i_ (Fig. 9A, Wilcoxon matched-pairs signed rank test, p=0.0156, n=6; Fig. 9C, p=0.0313, n=5). Further, we found a time delay between 5-HT activation of the epibranchial and nodose ganglia (Fig. 9B) comparable to the latency observed during hypoxia (Fig. 7F). Application of 5-HT_3_ receptor blocker, MDL72222, reduced the hypoxia-induced increase in [Ca^2+^]_i_ in the epibranchial ganglion (Fig. 9D, Wilcoxon matched-pairs signed rank test, p=0.0313, n=5). Further, 20 min incubation in tetrabenazine, a VMAT2 inhibitor, significantly reduced the increase in epibranchial [Ca^2+^]_i_ associated with hypoxia (Friedman test, p=0.0267, n=4), which fully recovered after 10 min washout (p=0.9590, n=4).

**Figure 9.**
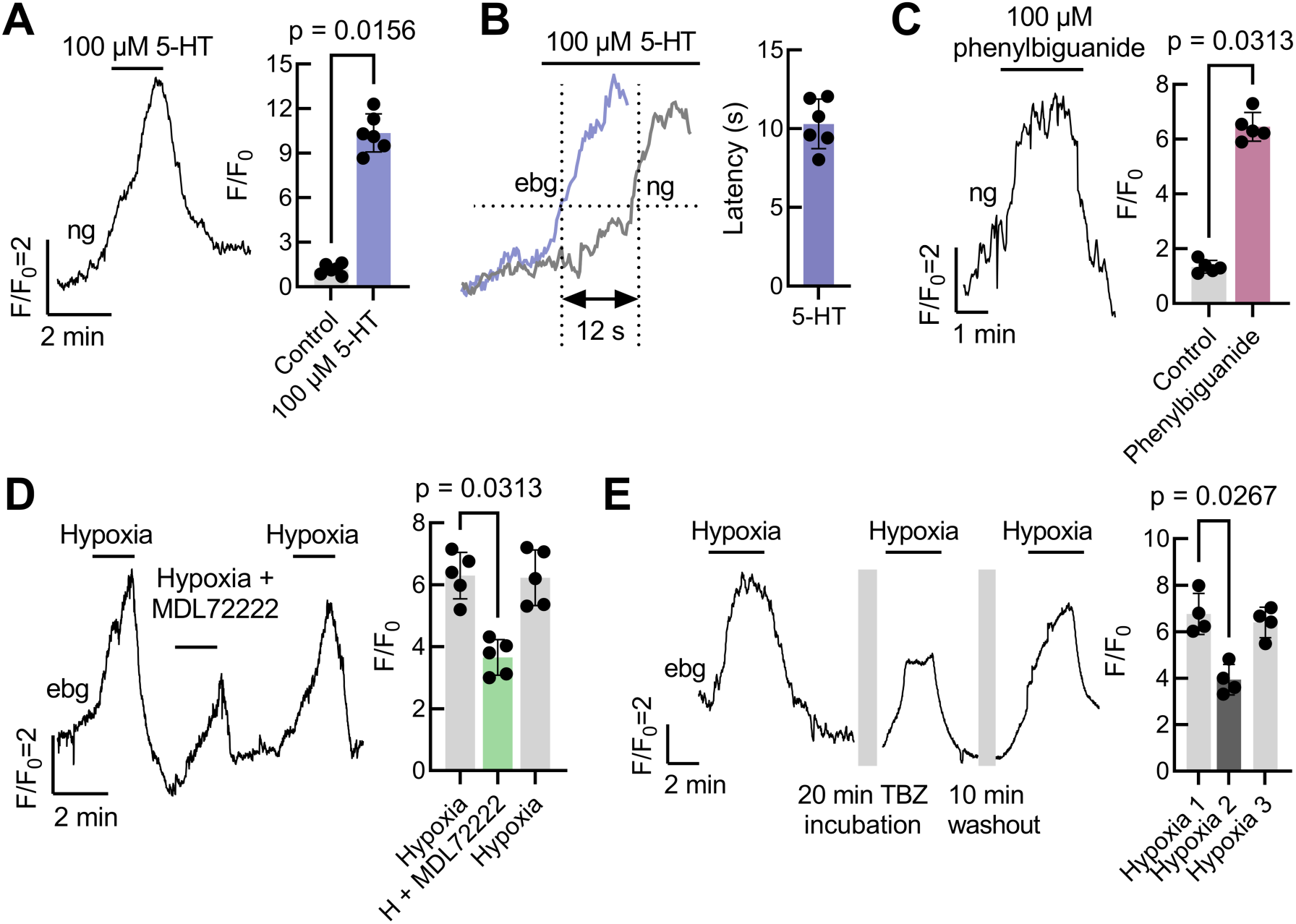
Vagal sensory ganglia increased intracellular calcium concentration ([Ca^2+^]_i_) in response to 5-HT_3_ receptor activation in whole-animal larval Ca^2+^ imaging. (A) Ca^2+^ imaging trace (left) and summary data with mean ± S.D. (right) from larval nodose ganglion (ng) responses to 5-HT. 5-HT induced an increase in [Ca^2+^]_i_ in this cluster of neurons (Wilcoxon matched-pairs signed rank test, p=0.0156, n=6). (B) Simultaneous Ca^2+^ imaging recording showing onset of the epibranchial (ebg) and nodose ganglion (ng) response to 5-HT (left). Onset of the [Ca^2+^]_i_ response was characterized as the time where relative fluorescence first doubled. At right, summary data showing mean ± SD time between onset of the epibranchial and nodose ganglia [Ca^2+^]_i_ responses to 5-HT. (C) Ca^2+^ imaging trace (left) and summary data with mean ± S.D. (right) from larval nodose ganglion (ng) responses to the 5-HT_3_ receptor agonist, phenylbiguanide. 5-HT_3_ receptor activation induced an increase in [Ca^2+^]_i_ (Wilcoxon matched-pairs signed rank test, p=0.0313, n=5). (D) Ca^2+^ imaging trace (left) and summary data with mean ± S.D. (right) from larval epibranchial ganglia in animals exposed first to hypoxia, followed by hypoxia in the presence of the 5-HT_3_ receptor antagonist, MDL72222. A third bout of hypoxia was applied to control for changes in fluorescence due to time. MDL72222 reduced the increase in epibranchial [Ca^2+^]_i_ associated with hypoxia (Wilcoxon matched-pairs signed rank test, p=0.0313, n=5). (E) Ca^2+^ imaging trace (left) and summary data with mean ± S.D. (right) from larval epibranchial ganglia before and after depletion of 5-HT by the VMAT2 blocker, tetrabenazine (TBZ). After an initial response to hypoxia was observed, the recording was paused (first break in trace, grey bar) for a 20 min incubation of the whole animal in 25 µM tetrabenazine. After TBZ incubation, a second bout of hypoxia was applied. The recording was paused during a 10 min washout period (second break in trace, grey bar), after which a third bout of hypoxia was applied to assess recovery. 20 min TBZ incubation significantly reduced the mean ± S.D. increase in [Ca^2+^]_i_ associated with hypoxia (Friedman test, p=0.0267, n=4), which was fully recovered after 10 min washout (Friedman test, p=0.9590, n=4).

Together, these results suggest the involvement of 5-HT excitatory neurotransmission in the gill acting via 5-HT_3_ receptors to mediate hypoxia signaling. However, in our isolated gill preparation, all concentrations of 5-HT tested failed to elicit a change in ChN [Ca^2+^]_i_ activity (Fig. 10A; Friedman test, p>0.9999, n=5), and application of MDL72222 had no effect on the ChN response to hypoxia (Fig. 10B, p=0.8125, n=5). Importantly, qPCR analysis confirmed expression of *htr3a* and *htr3b* (the genes encoding 5HT_3_ receptors) in the gill (Fig. 10C), though relative expression did not differ in gills from animals acclimated to hypoxia. Using confocal microscopy, we confirmed the location of 5-HT_3_ receptor activity in the gill. 5-HT_3_ receptor immunoreactivity revealed a punctuated labeling pattern that was confined to the distal region of gill filament (Fig. 10D1, white dashed box). At higher magnification and after rotation of image stacks, 5-HT_3_ receptor puncta were observed surrounding and in close association with S*-*type NECs (Fig, 10D2-7).

**Figure 10.**
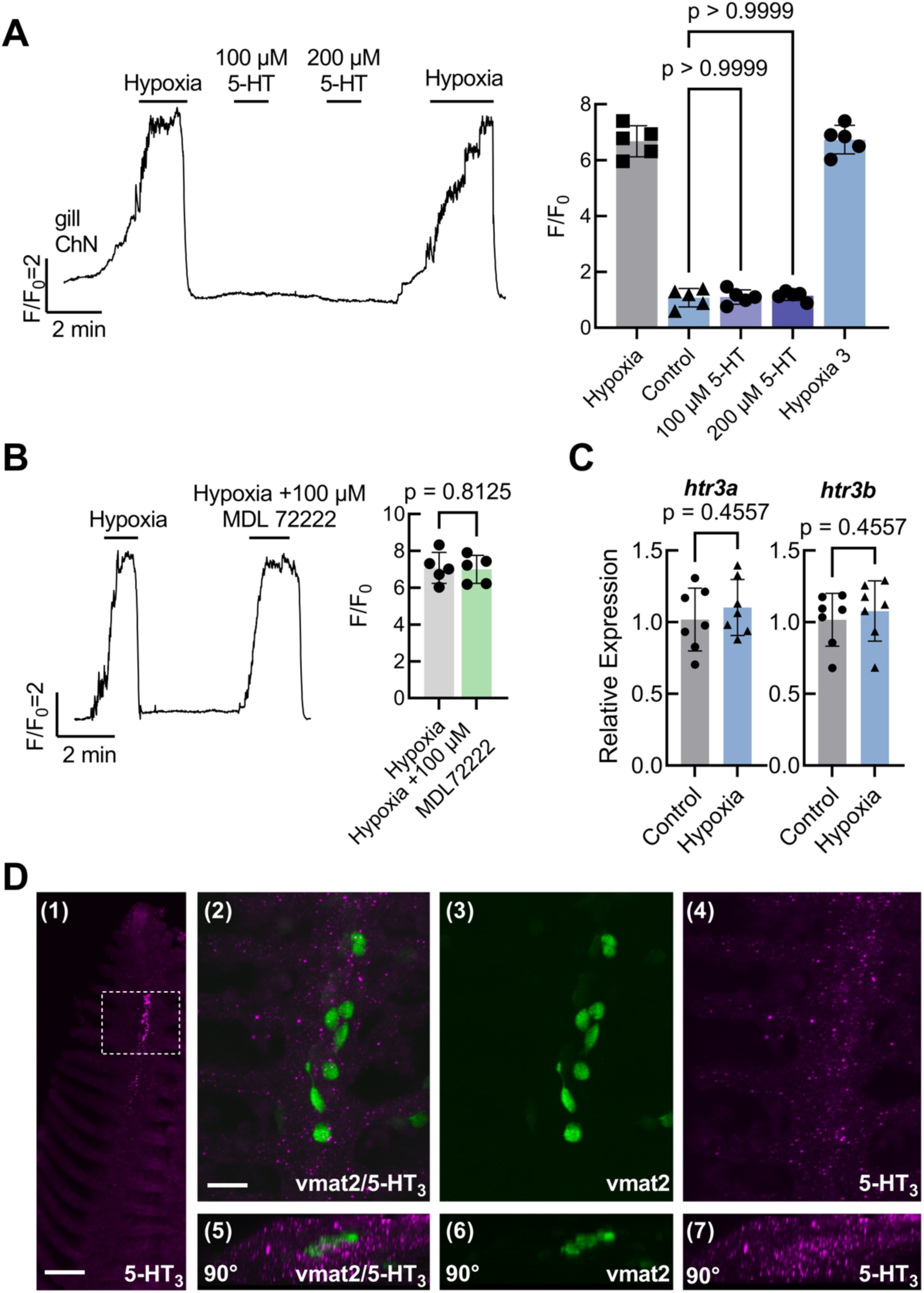
**Serotonergic hypoxia signaling occurred independently of the chain neuron pathway in the gill**. (A, left) Ca^2+^ imaging trace from a chain neuron (ChN) in an adult isolated zebrafish gill exposed to hypoxia followed by increasing concentrations of 5-HT (100 and 200 µM). A second bout of hypoxia was applied at the end of the recording to ensure viability of the preparation. (A, right) Summary data with mean ± S.D. from ChNs. Both concentrations of 5-HT tested failed to produce a Ca^2+^ signal significantly higher from the baseline control fluorescence (Friedman test, p > 0.9999, n=5). (B, left) Ca^2+^ imaging trace from a ChN in an adult isolated gill exposed to hypoxia followed by a second bout of hypoxia in the presence of the 5-HT_3_ receptor antagonist, MDL72222. (B, right) Summary data with mean ± S.D. from ChNs. 5-HT_3_ blockade did not affect the ChN Ca^2+^ response to hypoxia (Wilcoxon matched-pairs signed rank test, p=0.8125, n=5). (C) qPCR relative expression of *htr3a* and *htr3b* (genes encoding 5-HT_3_ receptor) in isolated gill tissue. Both genes were expressed in the gills with no changes in expression following 48 h of hypoxia exposure (Mann Whitney test, p=0.4557, n=7). (D1) 5-HT_3_ receptor immunoreactivity revealed a punctuated labeling pattern that was confined to the distal region of the gill filament (white dashed box). Scale bar in D1 = 50 µM. (D2) Immunolabeling of 5-HT_3_ receptors (magenta) revealed puncta that were often in close association with vmat2-positive neuroepithelial cells (NECs, green). Scale bar in D2 = 20 µM and applies to D3-7. (D3,4) *vmat2*-positive NECs and 5HT_3_ receptor labeling shown separately. (D5-7) Images from (D2-4) titled back 90° to show close association of 5-HT_3_ receptor labeling with *vmat2*-positive NECs.

## Discussion

This study provides evidence for dual excitatory pathways that mediate hypoxic signaling in the zebrafish gill. We previously demonstrated that GCaMP-positive gill NECs are oxygen-sensitive in *ex vivo* gill preparations (Reed and Jonz, 2025). Here, we further show that one chemoreceptor pathway includes a novel A-type NEC that stores ACh as a key excitatory neurotransmitter and activates postganglionic ChNs of the gill via nAChRs; and a separate pathway includes S-type NECs that store 5-HT and activate 5-HT_3_ receptors on terminals of neurons originating in the vagal sensory ganglia. Our study defines the functional organization and synaptic signaling mechanisms of oxygen chemoreceptors in zebrafish and is the first of its kind in a non-mammalian vertebrate model.

### Oxygen sensing via cholinergic excitation

The role of ACh in hypoxia sensing is well-characterized in the mammalian carotid body (Nurse, 2010; Kumar and Prabhakar, 2012; Leonard et al., 2018) but its role in fish gills has been controversial (Burleson and Milsom, 1995; Porteus et al., 2012; Shakarchi et al., 2013; Jonz et al., 2015). One challenge has been that chemosensory NECs in the zebrafish gill are routinely identified by labeling with 5-HT markers or expression of the monoamine transporter, *vmat2* (Jonz and Nurse, 2003; Pan et al. 2021). Non-serotonergic, NEC-like cells containing the vesicular ACh transporter, VAChT, were described in zebrafish (Shakarchi et al., 2013; Zachar et al., 2017b) and in an amphibious fish (*Kryptolebias marmoratus*; Regan et al., 2011), but the oxygen sensitivity of this cell type was never determined. In the present study, the localization of ACh immunoreactivity to SV2-positive NECs, but not to *vmat2*-expressing (serotonergic) NECs, confirmed the existence of a new subpopulation of neurosecretory cells, A-type NECs, that retain ACh but not 5-HT (summarized in Fig. 11).

**Figure 11.**
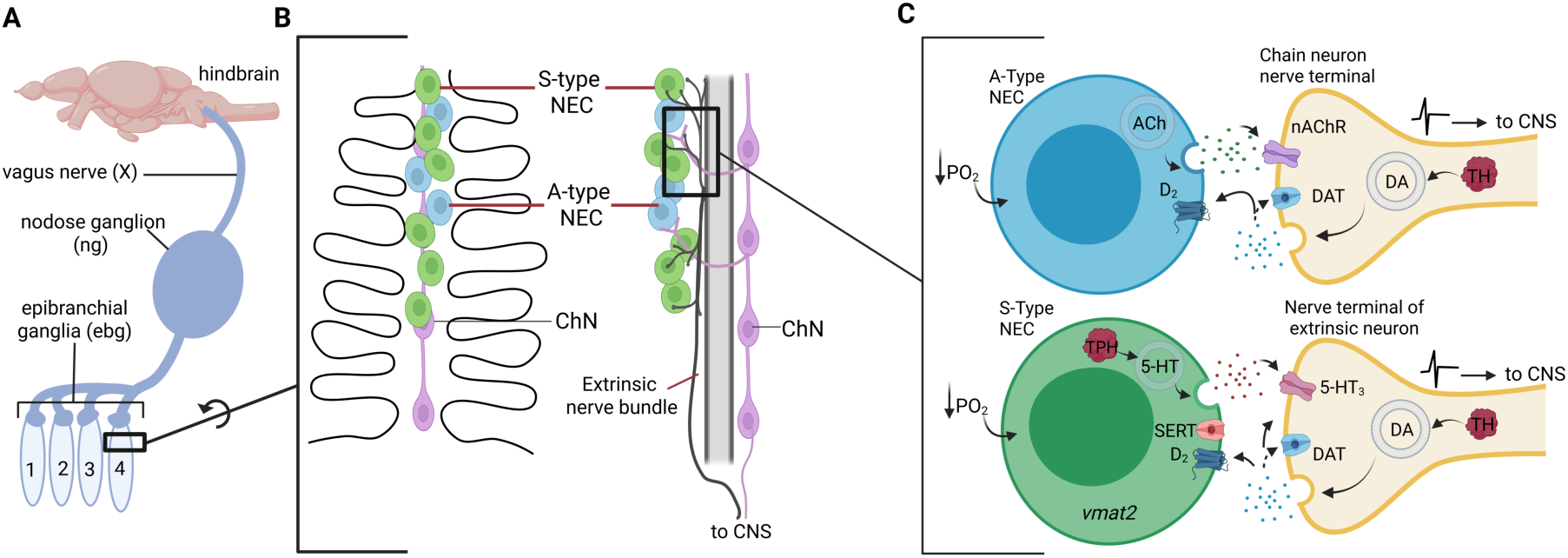
Schematic representation of a proposed mechanism of hypoxia signaling via two separate excitatory pathways: acetylcholine-type (A-type) NECs and chain neurons (ChN), and transmission via serotonin-type (S-type) NECs. (A) Schematic of cranial nerve X innervation of the gills, including the large vagus nerve, nodose ganglion (ng), and four epibranchial ganglia (ebg, 1-4) above the gill arches in a larval zebrafish. (B, left) highlighted view of a single gill filament showing distribution of A-type NECs (blue), S-type NECs (green), and postsynaptic chain neurons (ChN, magenta). (B, right) Vertical rotation of gill filament by 90° to show innervation of A-type NECs by ChN projections (magenta), which travel around the filament artery (center); and of S-type NECs by nerve terminals from the extrinsic nerve bundle (grey), which originates from neuronal cell bodies in vagal sensory ganglia. Our model suggests that ChNs act as interneurons only in the A-type NEC pathway, and both S-type NEC and A-type NEC pathways convergence or run in parallel before exiting the gill. (C) Close up view of A-type and S-type NEC signaling. Postsynaptic terminals for both pathways are shown in yellow. A-type NECs contain acetylcholine (ACh) and transmit the hypoxia stimulus via nicotinic receptors (nAChR) on postsynaptic ChNs. S-type NECs contain serotonin (5-HT) and transmit the hypoxia stimulus via 5-HT_3_ receptors of postsynaptic nerve terminals associated with ganglionic neurons. Our model assumes that both A-type and S-type NEC responses to hypoxia are modulated by dopamine (DA). This includes postsynaptic synthesis by tyrosine hydroxylase (TH), release of DA and uptake via the dopamine active transporter (DAT), and acting via presynaptic dopamine D_2_ receptors (Reed and Jonz, 2025; Reed et al., 2024). S-type NECs express tryptophan hydroxylase (TPH) for 5-HT synthesis, the vesicular monoamine transporter (VMAT2) and the serotonin transporter (SERT) (Pan et al., 2021, 2022).

In the present investigation, exogenous application of ACh, nicotine and hexamethonium effectively led to characterization of nAChRs on postsynaptic ChNs as a critical component of this oxygen-sensing pathway. A key observation in our study was the enhancement of ChN [Ca^2+^]_i_ responses to nicotine during hypoxia until a saturating concentration above 100 μM, where the additive effect was occluded. This suggested that hypoxia and nicotine were both acting through the same postsynaptic pathway. Moreover, the ChN [Ca^2+^]_i_ response to hypoxia, which must have occurred following endogenous release of a neurotransmitter, was progressively attenuated with increasing concentrations of hexamethonium. Since ChNs do not respond directly to hypoxia (Reed and Jonz, 2025), our findings strongly indicate that hypoxia triggers secretion of ACh from A-type NECs, and this causes activation of nAChRs on postsynaptic ChNs.

Evidence confirming the presence of nAChRs in the gill was lacking. Our results demonstrate expression of several nAChR subunits, including α2, α3, α4, α6, α7, β2, β3, and β4. Notably, acclimation to hypoxia for 48 h upregulated expression of α2, α3, β2, and β4 subunits, suggesting their specific involvement in the adaptive response to hypoxic environments. The upregulation of nAChR subunits in the gill in response to hypoxia bears some similarities to the carotid body, where chronic hypoxia induces upregulation of nAChR subunits (α3 and α7) on petrosal neurons that provide afferent innervation to the carotid body (Dinger et al., 2003). He et al. (2005) extended those findings by showing increased nAChR-mediated responses in afferent fibers that innervate type 1 cells.

In zebrafish, previous RNA-sequencing studies implicated α2 and β4 subunits specifically in gill neurons (Pan et al., 2022). The α2 subunit can form heteromeric receptors with either β2 or β4 subunits, conformations that are known to mediate excitatory neurotransmission (Whiteaker et al., 2009). Together, these results point to the potential formation of functional nAChRs in gill neurons, likely involving combinations of α2, β2, and/or β4 subunits that contribute to ChN activation during hypoxia. Our immunohistochemical data confirm that α2 subunit immunoreactivity was localized to zn-12-positive nerve fibers that were intimately associated with A-type NECs, and which were found surrounding the efferent filament artery. ChN cell bodies are located beneath the filament artery and are prime candidates to innervate NECs (Jonz & Nurse, 2003), suggesting that the α2-positive nerve fibers we observed near NECs are terminals of ChNs (summarized in Fig. 11).

Our study further investigated the potential role of ACh in respiratory control in zebrafish by exploring the involvement of cholinergic signaling in cranial nerve ganglia, which carry afferent signals required to initiate reflex hyperventilation. Using transgenic larvae expressing GCaMP6s, we examined changes in [Ca^2+^]_i_ in vagal sensory ganglia (nodose ganglion and epibranchial ganglia) in response to nicotine and ACh. The nodose ganglion responded to both ACh and nicotine in a manner similar to its response during hypoxia, suggesting that cholinergic signaling arising from ChNs in the gill is transmitted towards respiratory control centers in the hindbrain. Further, a significant time delay was observed between activation of the epibranchial ganglia (directly above the gill arches) and the nodose ganglion. This latency in activation indicated directional, time-dependent transmission of the hypoxic signal away from the gills. Additionally, our larval preparation showed responses to hypoxia in all four epibranchial ganglia, demonstrating that all four gill arches respond to hypoxia. These data support the idea that ACh released from A-type NECs in the gill mediate signaling to higher-order centers in the brain that control the hyperventilatory response to hypoxia.

### Oxygen sensing via serotonergic excitation

In parallel with the cholinergic pathway, we demonstrate that the sensing of hypoxia in the gill occurs through a separate pathway involving 5-HT. Serotonin has long been implicated as a neurotransmitter in gill NECs (Dunel-Erb et al., 1982; Burleson and Milsom, 1995; Jonz and Nurse, 2003; Porteus et al., 2012). In whole-animal experiments in zebrafish, hyperventilation was induced by 5-HT and the hyperventilatory response to hypoxia could be attenuated by MDL72222, a specific antagonist of ionotropic 5-HT_3_ receptors (Shakarchi et al., 2013; Jonz et al., 2015). In the present study, we discovered that application of 5-HT evoked time-dependent [Ca^2+^]_i_ activity in the epibranchial ganglia followed by the nodose ganglion. Stimulation in the nodose ganglion was reproduced by administration of the 5-HT_3_ receptor agonist, phenylbiguanide, and hypoxia-induced activity in the epibranchial ganglia was attenuated by coapplication of MDL72222. Critically, we did not find any 5-HT-dependent activity in ChNs. Instead, our experiments with the VMAT2 inhibitor, tetrabenazine, which reduced the hypoxic [Ca^2+^]_i_ response in epibranchial ganglia, suggest that S-type NECs in the gill initiate 5-HT-dependent activation of vagal sensory ganglia. Serotonergic NECs express *vmat2* for vesicular storage of 5-HT (Pan et al., 2021, 2022).

As with other receptor families, information on 5-HT receptor expression in the gill is incomplete (Reed and Jonz, 2022), though recent RNA-sequencing identified high expression of a 5-HT_3A_-like receptor subunit in gill neurons in zebrafish (Pan et al., 2022). In the present study, we confirmed expression of *htr3a* and *htr3b*, both of which encode subunits to form 5-HT_3_ receptors. Using immunohistochemistry, we further placed expression of the 5-HT_3B_ subunit to nerve terminals surrounding S-type NECs, the latter of which were labeled by *vmat2* expression. Given that no postganglionic neurons intrinsic to the gill were activated during 5-HT stimulation, our results lead us to conclude that S-type NECs are innervated directly by neurons of the epibranchial ganglia, which express postsynaptic 5-HT_3_ receptors and mediate hypoxic signaling to the hindbrain.

### Physiological and evolutionary implications

The existence of two distinct populations of oxygen chemoreceptors in fish gills, A-type and S-type NECs, suggests a more complex hypoxia signaling system than previously thought. The location of both NEC types in the gill epithelium, near the filament artery and external environment, puts them in an ideal position for sensing changes in environmental and/or arterial PO_2_. By comparison with mammalian chemoreceptors, one plausible explanation for this arrangement is that S-type NECs are more attuned to changes in external PO_2_, similar to serotonergic NEBs that mediate oxygen sensing in the airways of neonatal mammals (Cutz et al., 1993); whereas A-type NECs may detect arterial PO_2_, like type 1 cells in the mature carotid body (Ortega-Sáenz and López-Barneo, 2020). Evidence from developing zebrafish supports this idea. Gill NECs are first functional at 7 days post-fertilization, a time when 5-HT is involved in hypoxic responses and zebrafish rely primarily on diffusional gas exchange (Pelster and Burggren, 1996; Rombough, 2002; Jonz and Nurse, 2005; Shakarchi et al., 2013). Adult-like, ACh-dependent ventilatory reflexes, and NECs that express VAChT, do not appear until later developmental stages, when larvae are larger and depend on delivery of oxygen from the gills via the circulatory system (Jacob et al., 2002; Shakarchi et al., 2013).

Due to the homology between the carotid body and the first gill arch in fish, NECs were believed to be homologues of type 1 cells, the latter of which are derived from the embryonic neural crest (Pearse et al., 1973; Milsom and Burleson, 2007). Recent evidence has suggested that NECs containing 5-HT arise from the endoderm and may have a closer evolutionary link to pulmonary NEBs (Hockman et al., 2017), although single-cell RNA-sequencing demonstrated that gill NECs isolated from zebrafish express the neural crest marker, *ascl1* (Pan et al., 2022), a gene homologous to *Mash1* that is required for the formation of carotid body type I cells in mouse (Kameda, 2005). Taken together, previous investigations combined with the present results suggest the following. S-type NECs are endoderm-derived homologues of NEBs, are the first to mediate oxygen sensing in the gills, and may be external sensors of PO_2_. A-type NECs, on the other hand, contribute later to oxygen-sensing responses, when the gills are fully developed, and their morphological and physiology features suggest homology with type I cells that detect arterial PO_2_.

An important observation from the present study was that the relative expression of genes encoding subunits of nAChRs increased in zebrafish exposed to hypoxia for 48 h, whereas relative expression of 5-HT_3_ receptors did not. These findings raise the possibility that nAChRs and the A-type NEC pathway may also play a role in acclimatization to hypoxia. Fish naturally experience chronic episodes of hypoxia in their environment, which can lead to an increase in sensitivity to acute hypoxia, including increased hyperventilatory responses (Perry et al., 2009).

## Conclusion

This study provides new insights into the complex signaling mechanisms underlying oxygen sensing in the zebrafish gill. We identify two distinct populations of chemoreceptors that contribute to hypoxia signaling by independent postsynaptic mechanisms. We demonstrate that ACh is a key excitatory neurotransmitter involved in the activation of postganglionic neurons during hypoxia, whereas 5-HT acts through extrinsic innervation to the gill. Both pathways activate vagal sensory ganglia and may contribute to reflex hyperventilation. We argue that zebrafish possess oxygen chemoreceptors that are homologous with NEBs and type I cells, and provide a blueprint for understanding the diversity of oxygen sensing mechanisms and how oxygen sensing may have evolved in vertebrates.

## Acknowledgements

The authors wish to thank Dr. Tuan Bui for assistance with the GCaMP transgenic line during the early stages of this work. Funding was provided by the Natural Sciences and Engineering Research Council of Canada grant nos. 2018-05571 and 2024-03908.

